# Mesenchymal *Igf2* is a major paracrine regulator of pancreatic growth and function

**DOI:** 10.1101/714121

**Authors:** Constanze M. Hammerle, Ionel Sandovici, Gemma V. Brierley, Nicola M. Smith, Warren E. Zimmer, Ilona Zvetkova, Haydn M. Prosser, Yoichi Sekita, Brian Y.H. Lam, Marcella Ma, Wendy N. Cooper, Antonio Vidal-Puig, Susan E. Ozanne, Gema Medina-Gómez, Miguel Constância

## Abstract

The genetic mechanisms that determine the size of the adult pancreas are poorly understood. Here we demonstrate that many imprinted genes are highly expressed in the pancreatic mesenchyme, and explore the role of *Igf2 in-vivo*. Mesenchyme-specific *Igf2* deletion results in acinar and beta-cell hypoplasia, postnatal whole-body growth restriction and maternal glucose intolerance during pregnancy. Surprisingly, mesenchymal mass is unaffected, suggesting that the mesenchyme is a developmental reservoir of IGF2 used for paracrine signalling. The unique actions of mesenchymal IGF2 are demonstrated by the absence of phenotypes upon *Igf2* deletion in the developing pancreatic epithelium. Furthermore, increased IGF2 activity specifically in the mesenchyme, through *Igf2* loss-of-imprinting or *Igf2r* deletion, leads to pancreatic acinar overgrowth. *Ex-vivo* exposure of primary acinar cells to exogenous IGF2 increases cell proliferation and amylase production through AKT signalling. We propose that mesenchymal *Igf2*, and perhaps other imprinted genes, are key developmental regulators of adult pancreas size and function.

## Introduction

The mammalian pancreas plays a central role in energy homeostasis, which is achieved by functionally and morphologically distinct exocrine and endocrine components. Optimal pancreatic function requires a match between pancreas size and the physiological demands of the host organism. However, the genetic determinants of organ size, and the mechanisms that achieve adequate relative organ size, are poorly understood. The size of the pancreas is thought to be fixed early in development, limited by the size of the progenitor cell pool that is set-aside in the developing pancreatic bud^1^. In addition to autonomous cues in the epithelium, pancreatic development is also dictated by non-autonomous signals from the mesenchyme. Mesenchymal cells overlie the developing pancreatic bud and provide critical signals for the expansion of both precursors and differentiated endocrine and exocrine cells^2^. These cells are present throughout pancreas organogenesis but the relative proportion of mesenchyme to epithelium shifts, with a dramatic reduction over time, as the epithelial cells expand. Recent elegant genetic manipulation approaches have shown that mesenchymal cells regulate pancreatic growth and branching at both early and late *in utero* developmental stages^2,3^. However, many of the mesenchymal signals that control these processes remain un-identified. Moreover, the factors required for cell fate decisions that specify individual pancreas cell type subsets are fairly well established but less is known about signalling pathways involved in proliferation and survival of cell types.

Insulin-like growth factors (IGF1 and IGF2) are small mitogenic polypeptides (^~^7KDa) with structural homology to pro-insulin. Although IGFs expression is ubiquitous in many cell types, they are most abundant in the cells and tissues of mesodermal origin, and form important components of stem cell niches^4,5,6,7,8,9^. *Igf2* transcription is regulated by genomic imprinting^10^, an epigenetic process that causes a subset of genes in the genome to be transcribed according to parental origin. Imprinted genes regulate key aspects of mammalian physiology, from growth to energy homeostasis^11^. In most mouse and human tissues, *Igf2* is only transcribed from the paternally inherited allele, with the maternal allele being repressed by a methylation sensitive CTCF-dependent boundary that restricts the access of downstream enhancers to the *Igf2* gene promoters^12^. In humans, reduced *Igf2* expression contributes to the intra-uterine growth restriction in patients with Silver-Russell syndrome^13^. Conversely, bialellic *Igf2* expression caused by loss of *Igf2* imprinting is observed in Beckwith-Wiedemann patients^14^, a syndrome characterized by somatic overgrowth, neonatal hypoglycaemia with variable penetrance and increased predisposition to tumours. In mice, increased supply of embryonic *Igf2* above biallelic expression can result in disproportionate overgrowth associated with heart enlargement, oedema and fetal death^15^. *Igf2* null mice are viable, depending on the genetic background, with almost half the adult body weight^15^.

Key questions about how IGF2-mediated growth effects are developmentally programmed and how IGF2 contributes to organ growth control *in-vivo* remain unanswered. The normal timing, location and duration of IGF2 supply at the organ level are likely to be crucially important. The pancreas serves as an interesting model for the study of these processes, as it requires tight spatial and temporal regulation of proliferation, differentiation and morphogenesis. Very little is known about the main sites of *Igf2* expression and imprinting during pancreas development. Moreover, most mouse transgenic studies conducted so far have been directed to the beta-cells, in which *Igf2* is overexpressed^16,17^ or knocked-out^18^, or focused on mice that overexpress *Igf2*^19,20^ or lack *Igf1* and *Igf2* constitutively^21^.

Here we describe the generation of a conditional knock-out allele for *Igf2* and perform systematic analyses of cell-type specific deletions that target the developing pancreatic mesenchyme (*Nkx3.2*-Cre), epithelium (*Ptf1a*-Cre) or beta-cell (*RIP*-Cre), with the overall aim of defining the paracrine and autocrine roles of IGF2 in pancreatic growth. We demonstrate that the mesenchyme is a developmental reservoir of IGF2 that acts as an essential signal for exocrine and endocrine pancreas growth. We found little evidence for autocrine or paracrine growth effects of IGF2 from the developing epithelium, which is consistent with the low levels of expression seen in the epithelium when compared to the mesenchyme. Our findings show that the pancreatic hypoplasia caused by *Igf2* deletion from the mesenchyme is associated with reduced whole-body weight around weaning and onwards. To our knowledge, this constitutes the first evidence that organ-specific *Igf2* expression in fetal life can program the growth trajectory of the entire organism in post-natal/adult life. Furthermore, we show that glucose homeostasis is impaired in pregnant females that lack pancreatic mesenchymal *Igf2*. Together, these results uncover previously unanticipated roles for mesenchymal *Igf2* in controlling the normal development and function of the pancreas, with important consequences for energy homeostasis regulation.

## Results

### Generation of *Igf2*^+/fl^ mice and pancreas cell-type specific *Igf2* knockouts

Mice containing a floxed *Igf2* allele (*Igf2*^+/fl^) were generated (see Methods and Supplementary Fig. 1) and crossed with transgenic strains that express Cre recombinase in the main cell types of the developing pancreas, *i.e. Nkx3.2*-Cre: mesenchyme; *Ptf1a*-Cre: epithelium, *i.e*. exocrine, endocrine and ducts; *RIP*-Cre: beta-cells. Recombination between the two *loxP* sites led to inactivation of the *Igf2* gene *via* deletion of all coding exons (Supplementary Fig. 2). To confirm the efficiency of the Cre transgenics, we used a recombinase inducible YFP reporter under the control of the *Rosa26* locus (*Rosa26YFP*-stop^fl/fl^). This reporter was bred into the *Igf2*^+/fl^ background to generate an *Igf2*^+/fl^; *Rosa26YFP*-stop^fl/fl^ strain. In these mice, Cre expression causes inactivation of *Igf2*, as well as expression of YFP, which can be identified by immunofluorescence and/or flow cytometry. Using these methods, we confirmed the specificity of each Cre line, showed that *Igf2* is inactivated efficiently and that *Igf2* is mainly expressed from the paternal allele in the diverse pancreatic cell types (Supplementary Figs. 3 and 4). All genetic crosses used throughout, unless otherwise stated, refer to paternal transmission of the *Igf2* floxed allele (*Igf2*^+/fl^) with maternal transmission of a Cre allele (*Nkx3.2*^Cre/+^; *Ptf1a*^Cre/+^ or *RIP*^Cre/+^).

### The mesenchyme is the main source of *Igf2* in the developing pancreas

We first sought to establish which major cell types within the developing mouse pancreas express *Igf2* transcripts and to determine their relative levels using fluorescence activated cell sorting (FACS). An expression timeline analysis, from E16 to adult, revealed that mesenchymal cells express the highest levels of *Igf2* mRNA when compared to beta-cells or non-mesenchyme (which is comprised of exocrine, endocrine, ductal and endothelial cells) (Fig. 1a). At E16, mesenchyme cells express 380 fold more *Igf2* than beta-cells and 1.5 fold more than non-mesenchyme cells. At P14, the difference between mesenchyme and non-mesenchyme is greatest, with mesenchymal cells expressing 700 fold more *Igf2* than non-mesenchyme cells. The levels of *Igf2* in mesenchyme remain high in the neonatal period until the weaning period (at P21 levels are 3.2 fold reduced compared to E16, followed by a steep decline in adulthood) (Fig. 1a). Interestingly, both non-mesenchyme cells and the beta-cells decrease their *Igf2* expression levels after E16. *In situ* hybridization analysis at P5 shows that *Igf2* transcripts are localized in mesenchymal structures surrounding vessels and ducts, with low to undetectable expression in beta and acinar cells (Fig. 1b).

**Figure 1:**
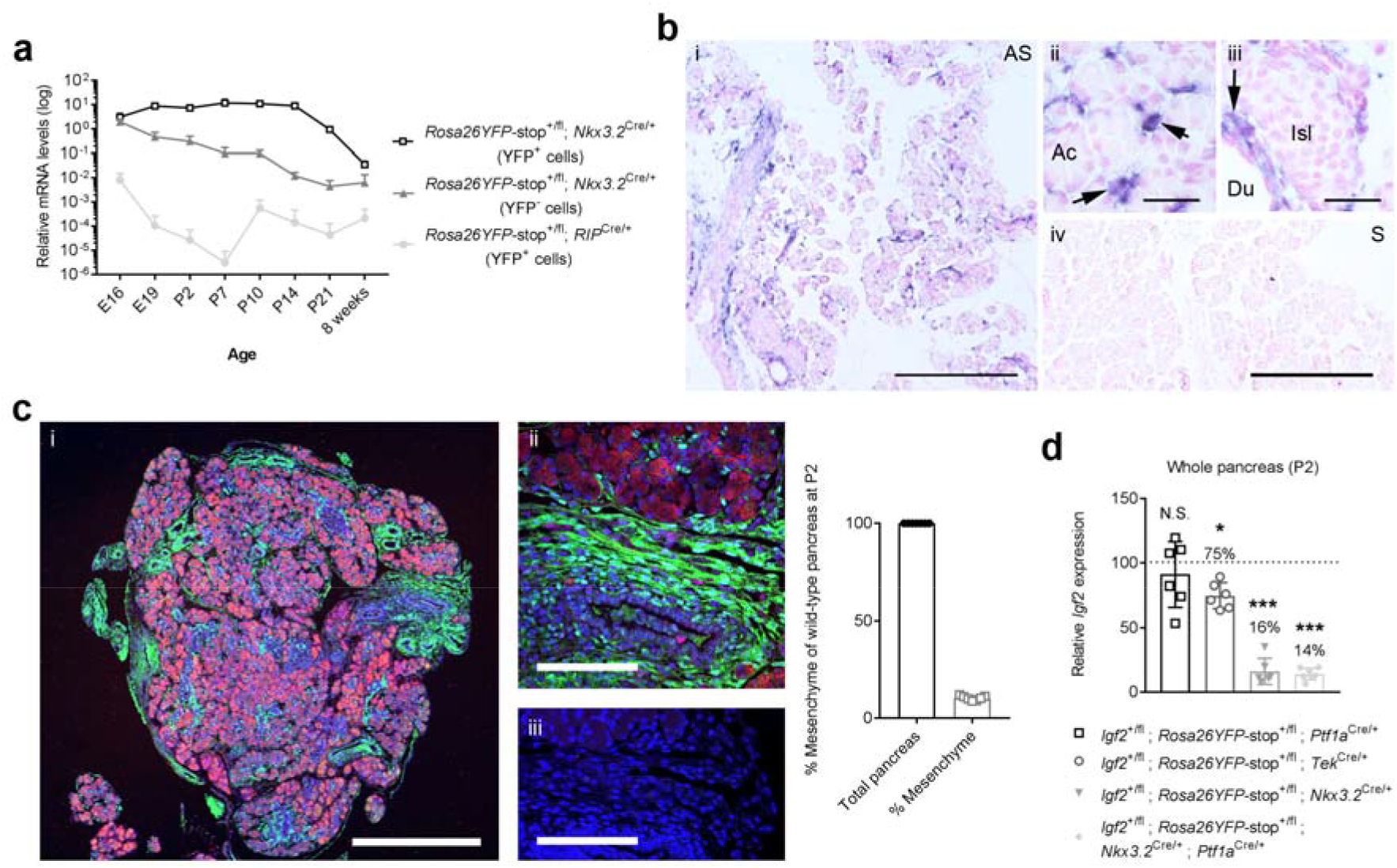
Developmental profiling of *Igf2* mRNA expression in mouse pancreas. (**a**) Timeline of *Igf2* mRNA expression measured by qRT-PCR in YFP+ or YFP–FACS isolated cells from offspring of *Nkx3.2*-Cre or *RIP*-Cre females mated with *Rosa26YFP*-stop^fl/fl^ males (E – embryonic day; P – postnatal day). Highest levels of *Igf2* expression are observed in the mesenchyme fraction (*Nkx3.2*-Cre YFP+ cells) throughout development. Expression data is normalized to *Ppia*, and is shown as average + SD (n=3–5 per time point and cell fraction). (**b**) *Igf2* mRNA analysis by *in situ* hybridization at P5 showing high levels of expression in mesenchymal cells (blue colour, see arrows). AS – antisense probe (in insets i, ii and iii); S – sense probe (negative control; in inset iv); Ac – acinar cells; Isl – pancreatic islet; Du – pancreatic duct. Scale bars: 30 μm and 300 μm in higher and lower resolution insets, respectively. (**c**) Representative immunofluorescence of a P2 pancreatic section (i) stained for acinar cell amylase (red), *Nkx3.2*-Cre driven YFP expression (green) as marker of mesenchyme cells and DAPI (blue) for nuclei. Scale bar: 0.5 mm. The insets show high levels of YFP in the connective tissue surrounding acinar cells (ii), and a negative control, without primary antibodies (iii). Scale bars for insets: 100 μm. The graph depicts the percentage of YFP+ pancreatic mesenchyme cells in wild-type pancreas (n=7). Data is shown as individual values. (d) Cell-type contribution to whole pancreas *Igf2* mRNA levels, measured by qRT-PCR in lysates at P2 from offspring of heterozygous Cre females mated with *Igf2*^+/fl^ males. Data was normalized to *Ppia* and shown as individual values and averages ± SD relative to levels in control littermates, which were arbitrarily set to 1. Residual levels of *Igf2* mRNA in each mutant are indicated as percentage values; (n=6-7 samples per genotype; N.S. – non-significant; * p<0.05; *** p<0.001 by unpaired Student’s *t*-test).

To measure more precisely the contribution of the various cell types to overall pancreatic *Igf2* levels, cell-type specific *Igf2* knockouts were analysed at the neonatal stage P2. The mesenchyme contributes approximately 10% of the pancreatic tissue at this time point, as measured by stereology (Fig. 1c). Notably, deletion of *Igf2* from those cells reduced total *Igf2* pancreatic levels to 16% of normal (Fig. 1d). *Ptf1a-Cre* mediated deletion of *Igf2* in exocrine, endocrine and ductal cells showed no discernible changes in whole pancreas *Igf2* mRNA, which is consistent with the *in situ* hybridisation data shown in Fig. 1b. Deletion of *Igf2* from endothelial cells (mediated by *Tek*-Cre) reduced *Igf2* mRNA to approximately 75-80% of control levels (Fig. 1d).

### Mesenchyme-specific *Igf2* signalling controls the growth of exocrine and endocrine pancreas, but has no autocrine actions

To assess whether mesenchymal *Igf2* plays a role in the programming of early growth of the pancreas, *Igf2*^+/fl^; *Nkx3.2*^Cre/+^ knockout mice were first analysed at the neonatal stage P2 (Fig. 2a,b). Mice with deletion of the paternal *Igf2* allele have normal body weight, but significantly lighter pancreases compared to littermate controls (69% of normal; Fig. 2c and Supplementary Fig. 5a). Deletion of the maternal *Igf2* allele has no phenotypic effects, which is consistent with findings that the maternal allele is transcriptionally inactive due to imprinting (Supplementary Figs. 5b,c). Importantly, *Nkx3.2-Cre* or *Igf2* floxed carrier mice are indistinguishable from wild-type littermates (Supplementary Figs. 5a,b). We next measured the number of cell nuclei in DAPI-stained paraffin sections to establish if the loss of pancreas weight was due to hypoplasia. We found that the number of cell nuclei was significantly reduced in mutants compared to controls (74% of normal – Fig. 2d). This finding suggests that mutant pancreases are smaller due a reduction in cell numbers, which is in agreement with the well-established role for IGF2 in the control of cell proliferation and survival. Stereological analysis revealed that loss of *Igf2* in pancreatic mesenchyme leads to decreases in acinar and beta cell mass, in line with the overall pancreas weight deficit (Fig. 2e). Also consistent with the decrease in acinar mass, mutant pancreases contained lower amounts of lipase (Fig. 2f). Interestingly, mesenchyme mass was not altered in mutants compared to controls, suggesting lack of autocrine actions of mesenchyme-derived IGF2 (Fig. 2e). To explore this hypothesis further, we performed a mesenchyme-specific deletion of the receptor that mediates the classic IGF2 actions on cell growth and survival^15^, the IGF type I receptor (IGF1R) (Supplementary Fig. 6a). Mutant pancreas weights at P2, as well as total body weights, were similar to littermate controls (Supplementary Fig. 6b). Thus, our data reveals that mesenchyme-derived IGF2 is likely to control acinar and beta cell growth through paracrine signalling from the mesenchyme, rather than via autocrine effects leading to loss of mesenchymal mass.

**Figure 2:**
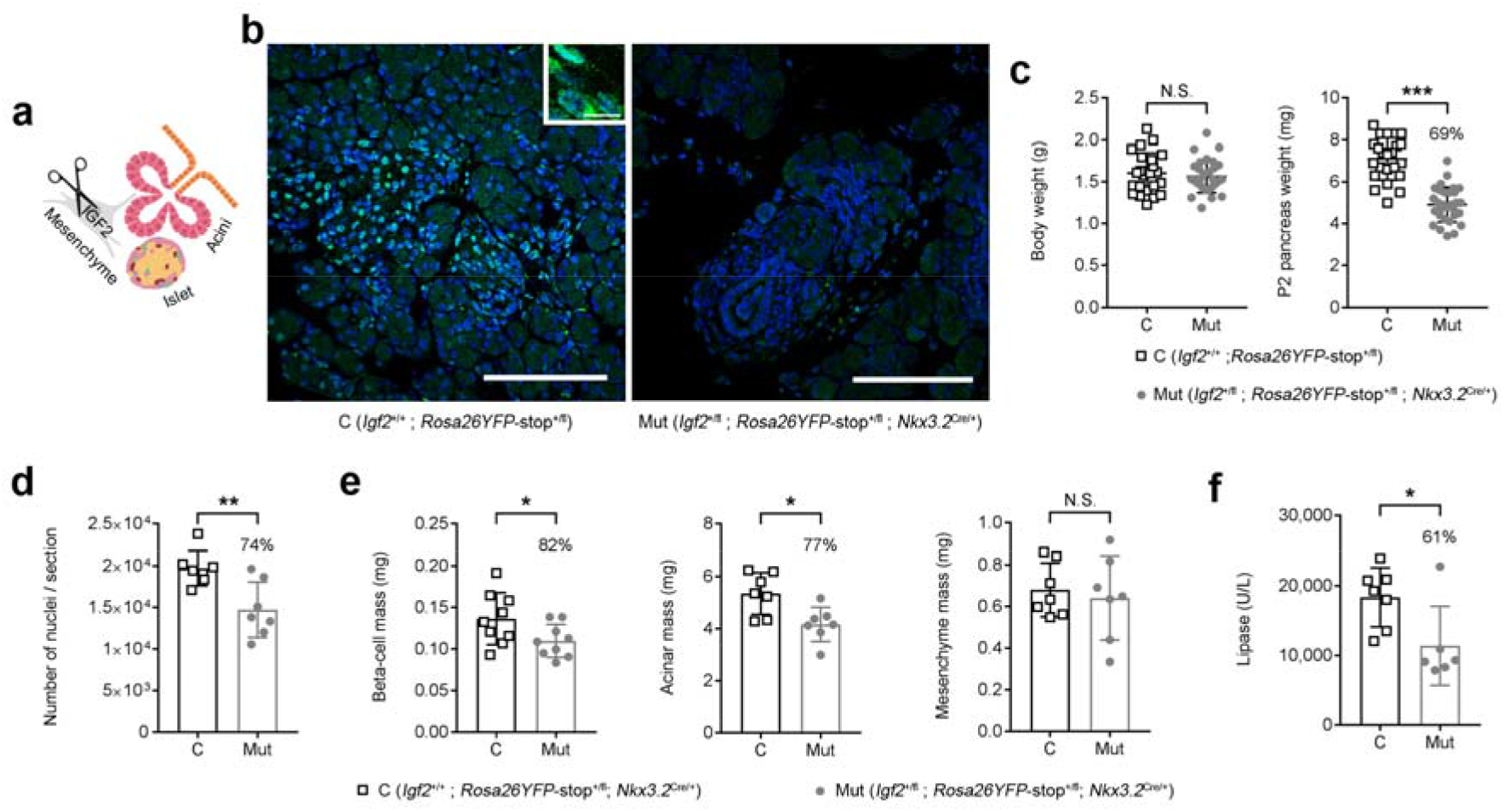
Pancreatic hypoplasia in mesenchyme-specific deletion of *Igf2* at postnatal day 2 (P2). (**a**) Schematic representation of conditional *Igf2* deletion from the pancreatic mesenchyme. (**b**) Representative immunofluorescence confocal microscopy showing cytoplasmatic staining (inset) for IGF2 in pancreata of control mice (C – *Igf2*^+/+^) and absence of signal in mutants (Mut – *Igf2*^+/fl^; *Nkx3.2*^Cre/+^) (green – IGF2; blue – DAPI staining of nuclei; scale bars – 100 μm and 10 μm for lower and higher magnification panels, respectively), (**c**) Pancreas weights are significantly reduced in mutants compared to littermate controls (n=23 controls and n=30 mutants), with similar body weights between genotypes, (d) Stereological measurements showing significant reductions (shown as %) in the total number of nuclei (measured in seven sections/sample). (**e**) Stereological measurements showing significant reductions in beta-cell mass and acinar mass, but not mesenchymal mass in mutants compared to littermate controls. (**f**) Reduced pancreatic lipase content in mutants compared to littermate controls. For all graphs, data is shown as average and individual values ± SD; N.S. – non-significant, * p<0.05, ** p<0.01 and *** p<0.001 by unpaired Student’s *t* tests.

### Growth effects caused by *Igf2* conditional inactivation are *miR-483* independent

The microRNA *miR-483* is located within the intron 4 of *Igf2*, which is deleted alongside the *Igf2* coding exons 4-6 upon Cre-mediated recombination. To rule out a contributory role of this microRNA to the phenotype, we analysed pancreases of mice carrying a *miR-483* deletion^22^, and found that it does not alter pancreas weight at P2 (Supplementary Fig. 7). Furthermore, *mir-483* knockout mice are normal sized, viable and show no evidence of impaired glucose homeostasis defects or other discernible phenotypes (Sekita *et al*. unpublished). We therefore conclude that the growth phenotype in the *Igf2*^+/fl^; *Nkx3.2*^Cre/+^ knockout mice can be attributed solely to IGF2 signalling actions.

### Mesenchyme-specific *Igf2 loss-of*-imprinting or *Igf2r* deletion result in pancreatic acinar overgrowth

We used a well-established mouse model of *Igf2* loss-of-imprinting – the H19DMD conditional knockout^23^ – to overexpress *Igf2* specifically in the pancreatic mesenchyme (*H19DMD*^fl/+^; *Nkx3.2*^+/Cre^) and investigated the effects on acinar growth (Fig. 3a). As expected, deletion of the imprinting control region (DMD) from the maternal allele resulted in increased *Igf2* and decreased *H19* mRNA levels (160% and 18% of controls, respectively – Fig. 3b) and was associated with a 29% increase in pancreas weight, with similar body weights between genotypes at P2 (Fig. 3c). The acinar cell mass was increased by 34%, in line with the overall pancreas weight increase (Fig. 3d). We then used an *Igf2* gain-of-function model that does not lead to transcriptional changes in *Igf2* or *H19* but instead results in increased IGF2 protein. Accordingly, we deleted the IGF-type II receptor that is required for recycling IGF2 via lysosome degradation^24^, specifically in the pancreatic mesenchyme (*Igf2r*^fl/+^; *Nkx3.2*^+/Cre^) (Fig. 3e). Reduced *Igf2r* mRNA levels in the mesenchyme cells (25% of controls – Fig. 3f), was associated with a 20% increase in pancreas weight, with similar body weights between genotypes at P2 (Fig. 3g). The acinar cell mass was also increased by 33% in this model (Fig. 3h). Therefore, the two gain-of-function genetic models further demonstrate that IGF2 produced by the pancreatic mesenchyme cells controls the growth of the acinar cells.

**Figure 3:**
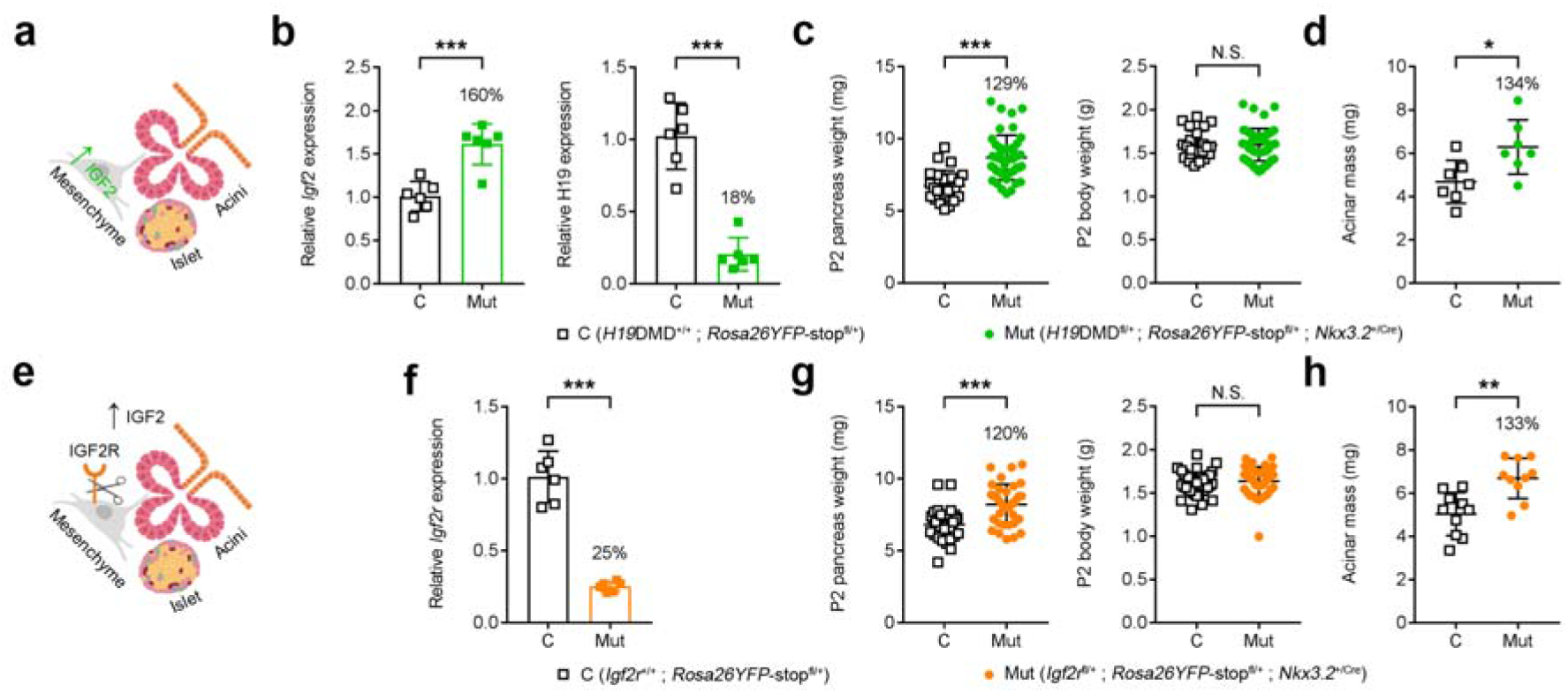
Pancreatic hyperplasia in two models of Igf2 gain-of-function, at postnatal day 2 (P2). (**a**) Schematic representation of conditional *Igf2* overexpression in pancreatic mesenchyme, (**b**) Increased *Igf2* mRNA expression and reduced *H19* mRNA levels in whole pancreata of mutant mice (Mut – *H19DMD*^fl/+^; *Nkx3.2*^+/Cre^) compared to control littermates (C – *H19DMD*^+/+^) measured by qRT-PCR. Data was normalized to *Ppia* and is shown relative to levels in controls (arbitrarily set to 1). (**c**) Pancreas weights, shown as individual measurements, are significantly increased in mutants compared to littermate controls (n=24 controls and n=45 mutants), with similar body weights between genotypes, (d) Stereological measurements revealed a significant increase (shown as %) in the acinar-cell mass in mutants compared to littermate controls. Data is shown as individual measurements. (**e**) Schematic representation of conditional *Igf2r* deletion in pancreatic mesenchyme, (**f**) Reduced *Igf2r* mRNA levels in whole pancreata of mutant mice (Mut – *Igf2r*^fl/+^; *Nkx3.2*^+/Cre^) compared to control littermates (C – *Igf2r*^+/+^) measured by qRT-PCR. Data was normalized to *Ppia* and is shown relative to levels in controls (arbitrarily set to 1). (**g**) Pancreas weights, shown as individual measurements, are significantly increased in mutants compared to littermate controls (n=39 controls and n=35 mutants), with similar body weights between genotypes, (**h**) Stereological measurements revealed a significant increase (shown as %) in the acinar-cell mass in mutants compared to littermate controls. Data is shown as individual measurements. For all graphs error bars represent SD, N.S. – non-significant, * p<0.05, ** p<0.01 and *** p<0.001 by unpaired Student’s *t* tests.

### The neonatal pancreatic mesenchyme is enriched in genes related to IGF signalling and imprinted genes

To investigate further the role of IGF2 signalling in the mesenchyme and its paracrine effects, we first performed genome wide-transcriptional profiling by RNA-seq in mesenchymal and non-mesenchymal cells isolated by FACS from P2 wild-type pancreata. 4,114 genes were found differentially expressed (fold change >1.5; FDR adjusted p value <0.05) between the mesenchyme and non-mesenchyme cells in wild-type *Igf2*^+/+^ pancreas (Supplementary Data 1). A number of these genes were validated by qRT-PCR in biological replicates (Supplementary Fig. 8a). Genes highly enriched in mesenchyme include several which have been shown to be expressed in embryonic pancreatic mesenchyme at E11.5^2,3^ (e.g. *Hgf, Sfrp1, Tgfb3, Tgfb2*) (Fig. 4a). Using DAVID functional annotation (see Methods) we identified over 20 significantly enriched GO terms (FDR adjusted p value <0.05) (Fig. 4b), which included biological processes known be involved in pancreatic mesenchyme function (e.g. Wnt signalling), early pancreas development (e.g. retinoic acid and SMAD signalling) and epithelial-mesenchymal interactions (integrin-mediated signalling, cell adhesion, collagen catabolism). This analysis also highlighted unexpected pathways, such as semaphorin-plexin and insulin-like growth factor receptor signalling (Fig. 4b). Several gene members of the IGF signalling family are highly enriched in mesenchyme versus non-mesenchyme (Fig. 4c), with *Igf2* being the highest expressed gene in pancreatic mesenchyme (Fig. 4d) and one of the most enriched compared to non-mesenchyme (53 fold; Fig. 4a and Supplementary Data 1). Interestingly, imprinted genes with known roles in growth control make up almost one third of the top 30 genes expressed in the pancreatic mesenchyme (Fig. 4d).

**Figure 4:**
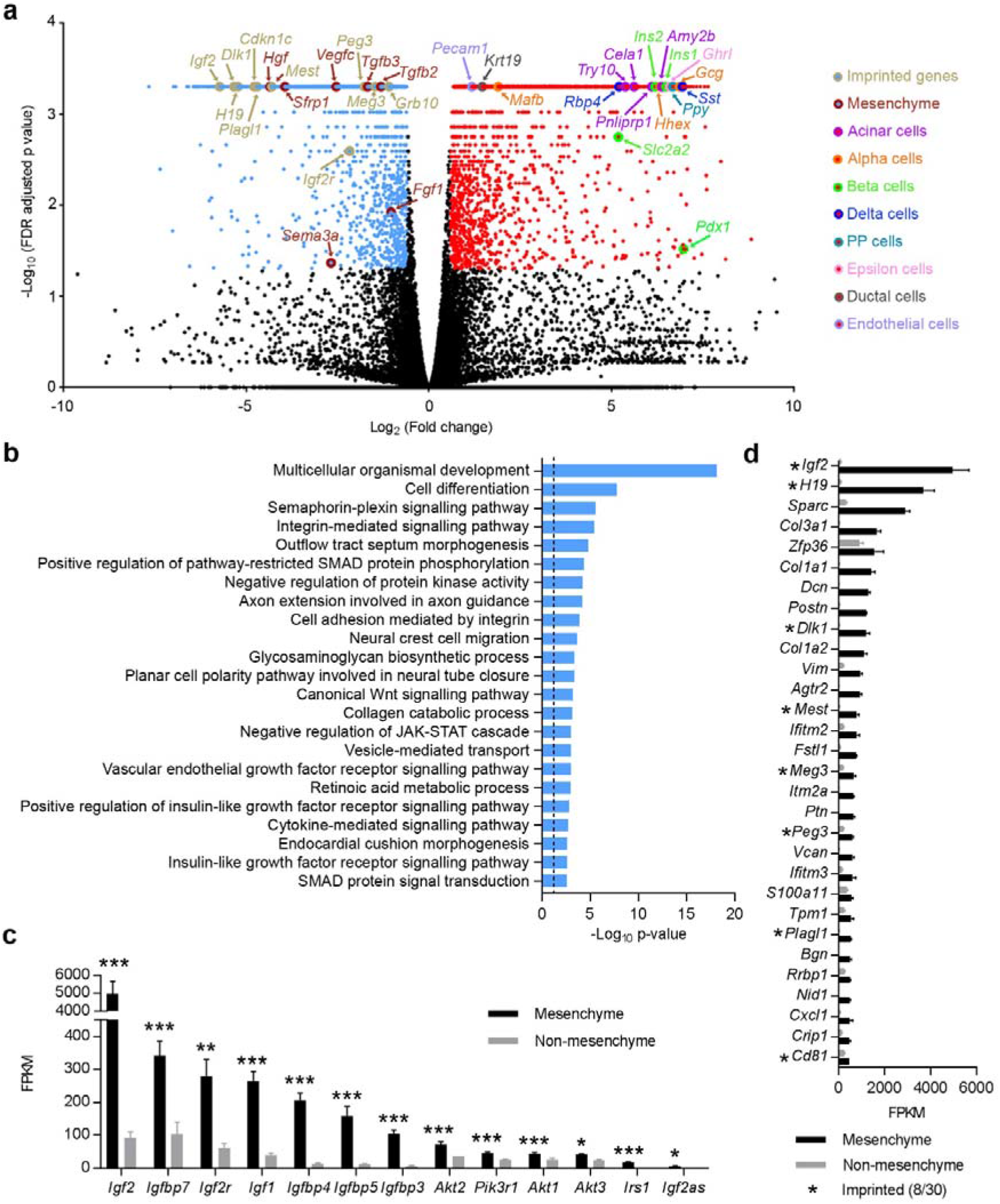
Distinct gene expression patterns of pancreatic mesenchyme identified by RNA-seq at postnatal day 2 (P2). (**a**) Volcano plot representing all expressed transcripts in pancreatic mesenchyme and non-mesenchyme cells (purified by FACS). For every transcript, the Log_2_ fold change between mesenchyme and non-mesenchyme cells was plotted against the −Log_10_ p value (FDR adjusted). Genes significantly enriched, with a fold change >1.5 and FDR adjusted p value <0.05, in mesenchyme are depicted as blue dots (n=1,902 genes), in non-mesenchyme as red dots (n=2,212), and those not significantly enriched in either cell fraction are depicted as black dots (all genes are listed in Supplementary Data 1). Selected genes of interest, such as known mesenchyme-expressed genes, signature genes for the various non-mesenchyme cell-types and imprinted genes are grouped by colour. (**b**) Top scoring biological processes containing genes enriched in the pancreatic mesenchyme as identified by DAVID functional annotation include IGF receptor signalling (ranked 19^th^ and 22^nd^). The enriched genes involved in this pathway are highlighted in (**c**). The dotted line in panel (**b**) corresponds to a p□value of 0.05. For panel (**c**) data is shown as average FPKM + SEM (n=4), ranked by levels of expression. FDR adjusted p values: * p<0.05, ** p<0.01 and *** p<0.001. (**d**) Top 30 expressed genes with highest FPKM values in pancreatic mesenchyme cells (average FPKM + SEM in mesenchyme and non-mesenchyme cells with n=4 per group). Note that all 30 genes are significantly enriched in mesenchyme compared to non-mesenchyme (>1.5 fold, FDR adjusted p value <0.05). Asterisks indicate imprinted genes, of which there are 8 amongst the top 30.

As expected, the non-mesenchyme *Igf2*^+/+^ fraction is highly enriched for genes expressed by different endocrine cell types, such as *Ins1* (90 fold enriched), *Pdx1* (126 fold), *Gcg* (116 fold) and *Sst* (123 fold) and genes encoding digestive enzyme such as *Amy2b* (83 fold), *Pnliprp1* (69 fold), *Cela1* (49 fold) and *Try10* (42 fold), (Fig. 4a and Supplementary Data 1). Supplementary Fig. 8b shows the top 30 expressed genes in the non-mesenchyme fraction.

### Mesenchymal *Igf2* loss-of-function leads to widespread transcriptional changes in the pancreatic epithelium

To investigate the molecular signatures associated with paracrine effects, we performed genome wide-transcriptional profiling by RNA-seq in both fractions of *Igf2*^+/fl^; *Nkx3.2*^Cre/+^ mutants (*i.e*. mesenchyme cells with *Igf2* deleted and the corresponding non-mesenchyme fraction), compared to mesenchyme and non-mesenchyme cells isolated from controls, at the neonatal P2 stage. We found that transcriptional changes are far more widespread in the non-mesenchyme fraction (498 differentially expressed genes with >1.5-fold change and FDR-adjusted p value <0.05) (Fig. 5a and Supplementary Data 2) than the *Igf2*-deficient mesenchyme itself (151 differentially expressed genes; Fig. 5a and Supplementary Data 3). DAVID functional annotation identified 15 enriched GO terms in the non-mesenchyme fraction (Fig. 5b and Supplementary Data 2) but only 2 in the *Igf2*-deficient mesenchyme fraction (namely, actin binding and inflammatory response, Supplementary Data 3). These findings are in agreement with the stereological analyses showing that loss of *Igf2* from the mesenchyme impacts on the growth/function of the non-mesenchymal fraction through a paracrine effect, but not the growth of the mesenchyme itself. The top biological processes in the non-mesenchyme fraction enriched in genes up-regulated in mutants were related to digestion, inflammation/immune responses, apoptosis and ERK1/2 signalling, and biological processes enriched in genes down-regulated in mutants were related to tRNA methylation and erythrocyte development (Fig. 5c and Supplementary Data 2).

**Figure 5:**
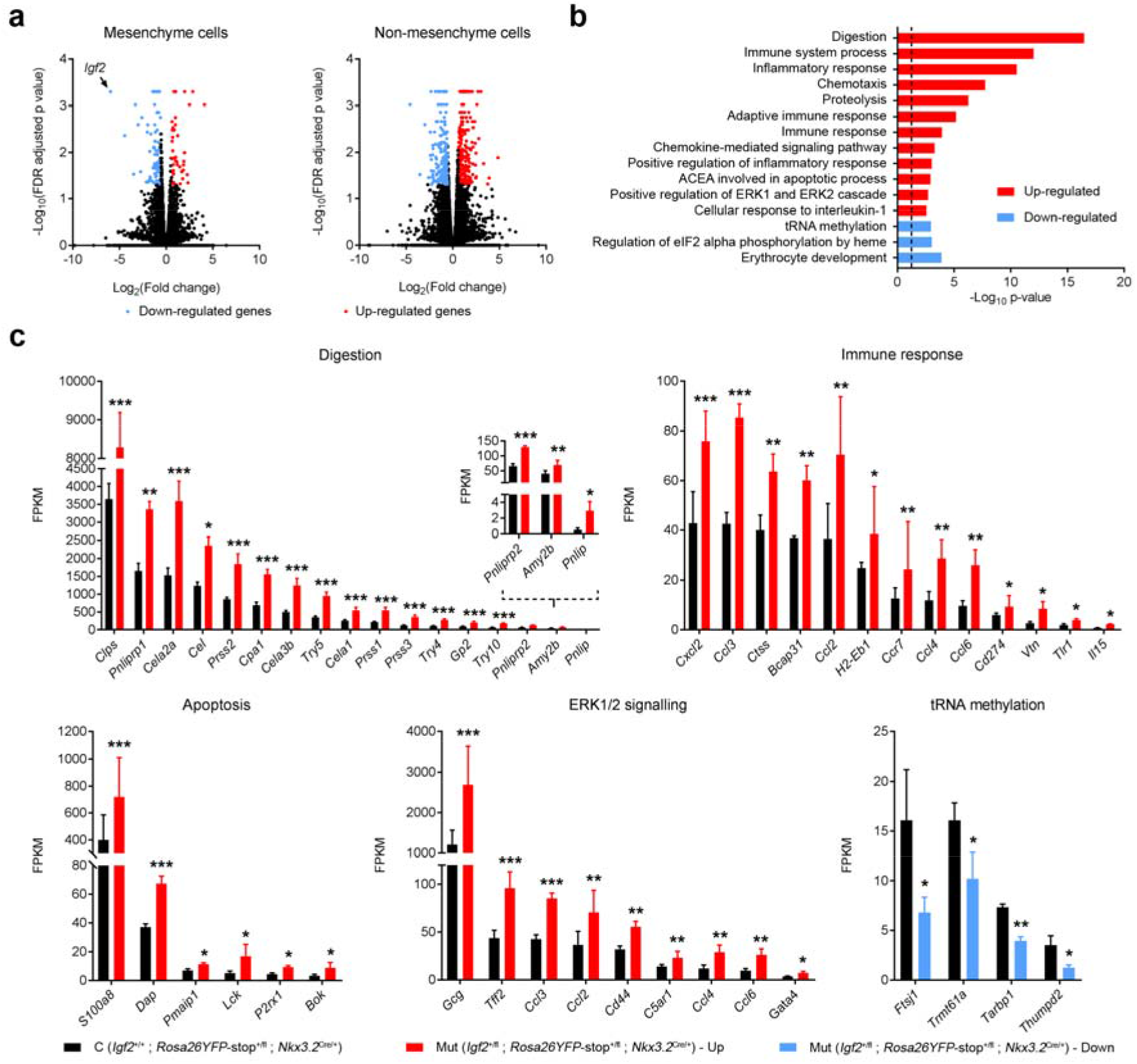
Loss of *Igf2* in pancreatic mesenchyme causes significant transcriptional changes in non-mesenchymal cells in a paracrine manner. (**a**) Volcano plot representing all expressed transcripts (by RNA-seq) in *Igf2*-deficient pancreatic mesenchyme (*Igf2*^+/fl^; *Nkx3.2*^Cre/+^) (left panel) and in corresponding non-mesenchyme cell fraction (right panel), compared to mesenchyme and nonmesenchyme controls (*Igf2*^+/+^; *Nkx3.2*^Cre/+^), respectively. Expression changes detected in the right panel thus reflect changes caused by lack of *Igf2* from the mesenchyme on neighbouring non-mesenchymal cells. For every transcript, the Log_2_ fold change of mutant versus control was plotted against the −Log_10_ p value (FDR adjusted). Blue and red dots depict statistically significant down-regulated and up-regulated genes (fold change >1.5 fold FDR adjusted p value <0.05), respectively, and black dots show genes that are not statistically different; (n=3 mutants and n=4 controls per cellular fraction). There are more differentially expressed genes in the non-mesenchyme fraction (216 down-regulated and 282 up-regulated genes) than in the mesenchyme (109 down-regulated and 42 up-regulated genes) (all genes are listed in Supplementary Data 2 and 3). (b) Biological processes enriched in down-regulated (blue) and upregulated (red) genes between nonmesenchyme of *Igf2*^+/fl^; *Nkx3.2*^Cre/+^ mutants and *Igf2*^+/+^; *Nkx3.2*^Cre/+^ controls as identified by DAVID. The horizontal axis shows −Log_10_ of p value; the dotted line corresponds to a FDR-corrected p□value of 0.05. (c) Differentially expressed genes in non-mesenchymal *Igf2*^+/fl^; *Nkx3.2*^Cre/+^ cells compared to *Igf2*^+/+^; *Nkx3.2*^Cre/+^ control cells, grouped according to selected enriched biological processes shown in b). Data is shown as average FPKM + SEM (n=4 *Igf2*^+/+^; *Nkx3.2*^Cre/+^ controls and n=3 *Igf2*^+/fl^; *Nkx3.2*^Cre/+^), ranked by levels of expression (the inset depicts lower expressed genes related to “digestion”, with a different scale bar). FDR adjusted p values: * p<0.05, ** p<0.01 and *** p<0.001.

### Mesenchymal cells secrete IGF2 and acinar cells respond functionally to exogenous IGF2 *ex-vivo*

To provide evidence that mesenchyme-derived IGF2 acts on acinar cells in a paracrine manner, we isolated primary mesenchyme and acinar cells from P2 pancreata (see Methods) (Fig. 6a,b). Mesenchymal cells secreted higher levels of IGF2 into the conditioned media compared to acinar cells (Fig. 6c), a finding that is consistent with higher levels of *Igf2* mRNA expression in the mesenchyme *in-vitro* and *in-vivo* (Fig. 6b and Fig. 1a,b). We then tested the effect of exogenous IGF2 treatment on acinar cells cultured *ex-vivo*. Isolated acinar cells treated with recombinant mouse IGF2 showed increasing levels of AKT phosphorylation (S473) in a concentration-dependent manner (Fig. 6d). Additionally, treatment of acinar cells with 50 ng/ml IGF2 stimulated their proliferation and led to a modest, but significant increase in amylase production (Fig. 6e). Altogether, the *ex-vivo* data shows that mesenchyme cells are capable of secreting IGF2 and that exogenous IGF2 induces intracellular signalling in acinar cells, associated with increased enzymatic output.

**Figure 6:**
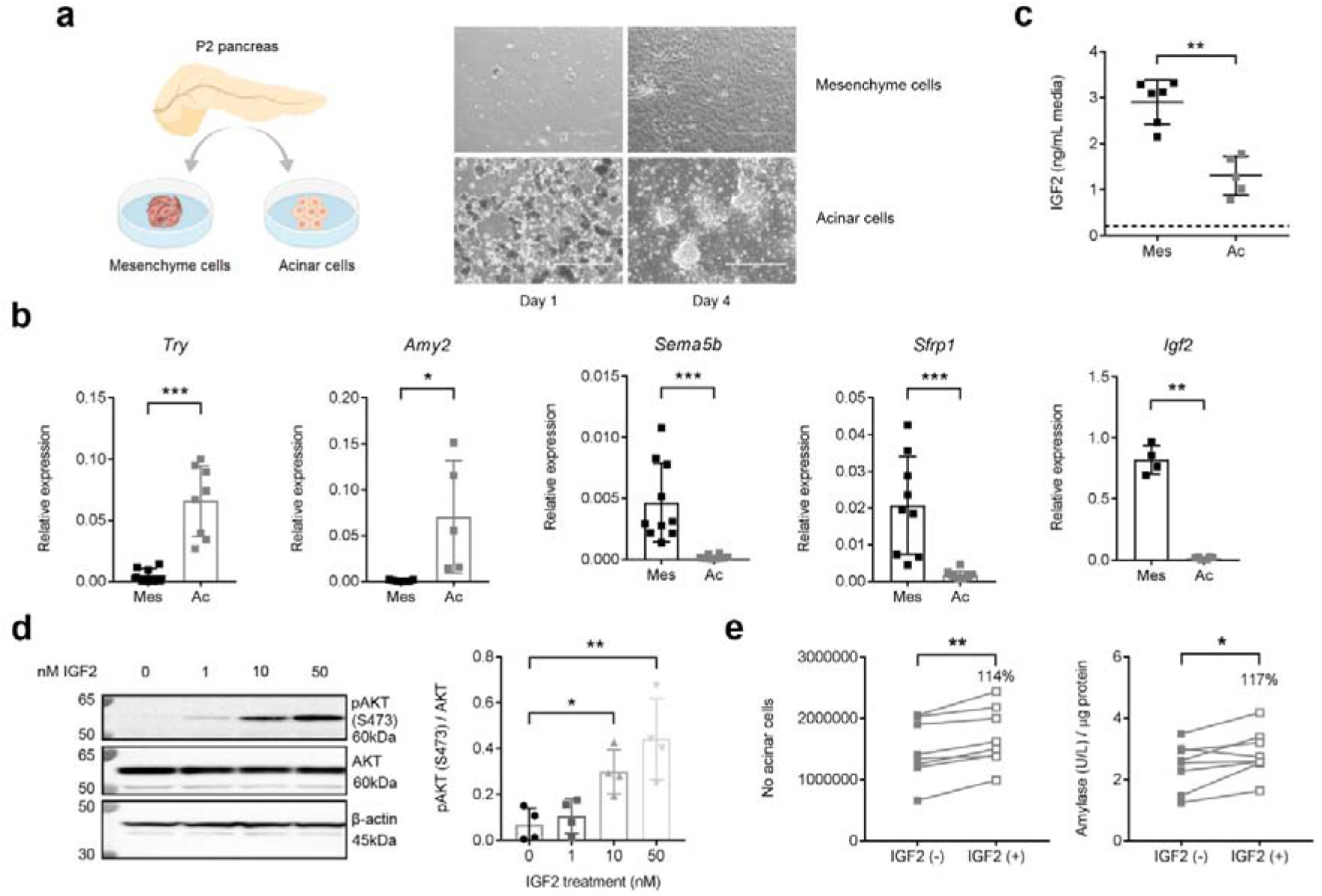
Paracrine effects of mesenchymal IGF2 on acinar cells using a primary cell culture model. (**a**) Primary mesenchymal (Mes) and acinar cells (Ac) were isolated from P2 pancreata. Representative images are shown after one and four days in culture. (**b**) mRNA expression levels of acinar (*Try, Amy2*) and mesenchymal (*Sema5b, Sfrp1, Igf2*) signature genes measured by qRT-PCR after four days of culture, showing high purity of the primary cell cultures, (**c**) Measurement of IGF2 protein secreted by mesenchymal or acinar cells in the culture media. Dotted line corresponds to background readings in media only (d) Levels of AKT phosphorylation (pAKT-S473) after 10 minutes treatment of freshly isolated acinar cells with increasing doses of IGF2 (0nM to 50nM) by western blotting, and quantified against total AKT levels (n=4 independent biological replicates). (**e**) Effect of IGF2 treatment (IGF2 +: 50 ng IGF2/ml) of primary acinar cells on total cell number (left panel) compared to non-treated cells [(IGF2 (-)], and amylase production normalized per protein content (right panel). For all graphs, the data is shown as individual values and averages, error bars represent SD. * p<0.05, ** p<0.01, *** p<0.001 by Mann-Whitney tests (**b, c**), Friedman’s test with Dunn’s correction for multiple comparisons (**d**) and the Wilcoxon matched-pairs signed rank test (**e**).

### Post-weaning growth and glucose homeostasis analyses in *Igf2* pancreas-cell type specific knockouts

To investigate the impact of *Igf2* loss-of-function from the developing mesenchyme and/or epithelium on post-weaning growth and glucose homeostasis, single and double *Nkx3.2*-Cre and *Ptf1a*-Cre *Igf2* knockouts, as well as beta-cell specific *RIP*-Cre *Igf2* knockouts, were analysed (Fig. 7a). The pancreas weight deficit observed in single *Igf2*^+/fl^; *Nkx3.2*^Cre/+^ knockouts at P2 is maintained at weaning (Supplementary Fig. 9a) and is also observed at 9 weeks of age in single *Igf2*^+/fl^; *Nkx3.2*^Cre/+^ knockout males and females (Fig. 7b). However, in contrast to the P2 time point, body weights are now reduced, by approximately 12% at P21 (Supplementary Fig. 9b), and by 5% and 10% for females and males, respectively, at 9 weeks of age (Fig. 7c). After normalization for body weight, the mutant pancreases remain disproportionately smaller at both P21 and 9 weeks time points (Supplementary Fig. 9a and Fig. 7b).

**Figure 7:**
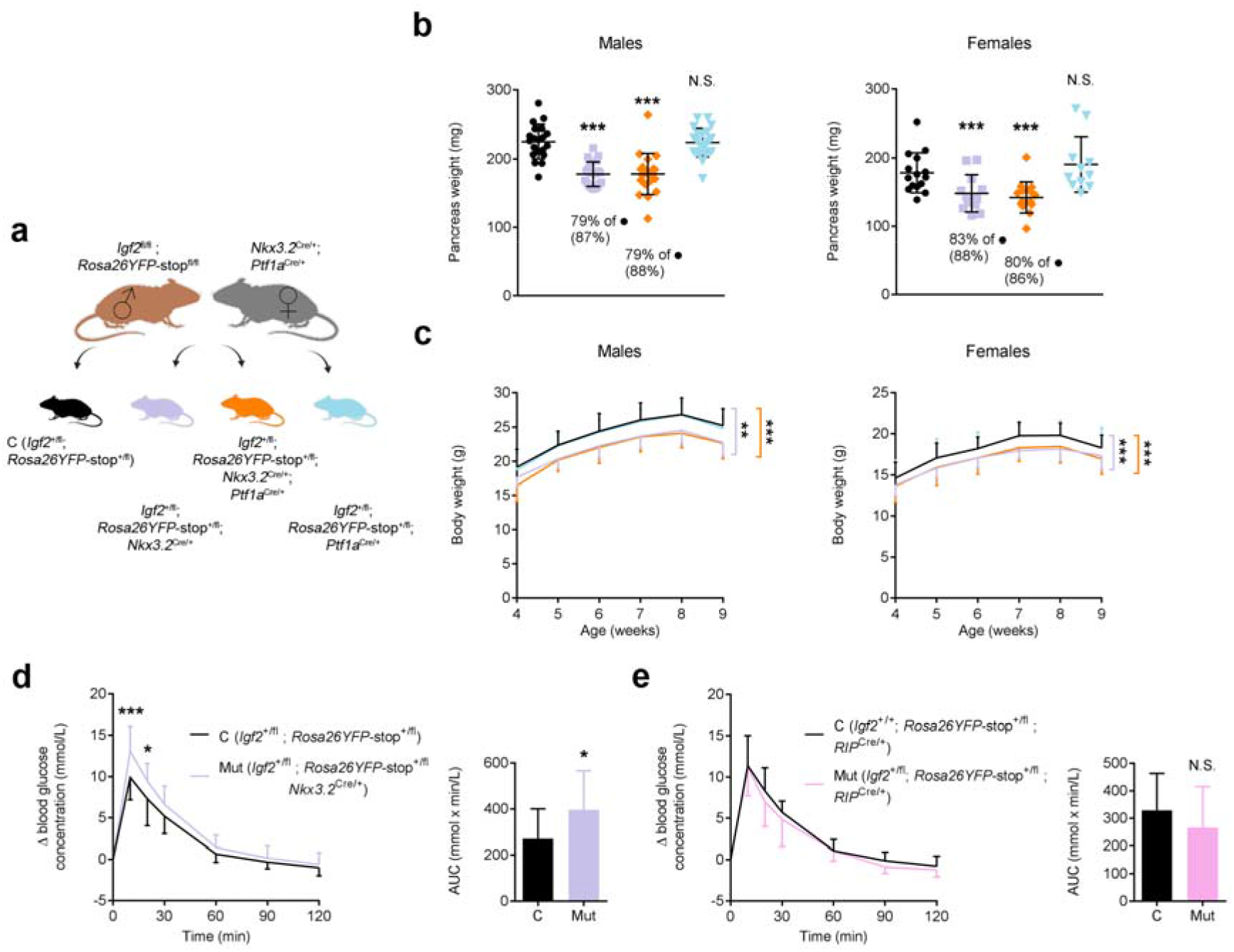
Growth and glucose homeostasis regulation in pancreas-cell type specific *Igf2* knockout mice in adulthood. (**a**) Schematic representation of the mating strategy used to generate control mice (black) and littermates with *Igf2* deletion in the pancreatic mesenchyme only (purple), mesenchyme plus epithelium (orange) or epithelium only (blue). (**b**) Pancreas weights at 9 weeks of age for males (n=20-22 per genotype) and females (n=11-15 per genotype). Significant reductions in weight (shown as %) are only observed in mice that carry a deletion of *Igf2* in the mesenchyme (i.e. *Igf2*^+/fl^; *Nkx3.2*^Cre/+^); values within brackets show % reduction after normalization to body weight; *** p<0.001 by 1-way ANOVA with Dunnett’s multiple comparisons tests against controls (C – *Igf2*^+/fl^). (**c**) Growth kinetics from weaning to 9 weeks of age for males (n=20-22 per genotype) and females (n=11-15 per genotype). ** p<0.01 and *** p<0.001 by repeated measures 1-way ANOVA with Dunnett’s post hoc test for multiple comparisons against the controls (C – *Igf2*^+/fl^). Oral glucose tolerance tests performed in pregnant 8-week old females at E15 of gestation after six hours of fasting: (**d**) – *Igf2* mesenchyme-specific deficient females (n=16 *Igf2^+/fl^* controls and n=15 *Igf2*^+/fl^; *Nkx3.2*^Cre/+^ mutants) and **(e)** *Igf2* beta-cell specific deficient females (n=15 *Igf2*^+/fl^, *RIP*^Cre/+^ controls and n=14 *Igf2*^+/fl^; *RIP*^Cre/+^ mutants). For both (**d**) and (**e**), changes in blood glucose concentrations (y-axis) from basal pre-treatment values with time (x-axis) after glucose administration are shown. * p<0.05 and *** p<0.001 by two-way ANOVA with Sidak’s multiple comparison tests. The graphs on the right side indicate area under curve (AUC) calculated using the trapezoid rule; * p<0.05 by unpaired Student’s *t* test. For all panels, data is shown as average values ± SD; N.S. – non-significant.

Double *Igf2*^+/fl^; *Nkx3.2*^Cre/+^; *Ptf1a*^Cre/+^ knockouts show similar pancreatic weights and body weight reductions to the single *Igf2*^+/fl^; *Nkx3.2*^Cre/+^ knockout (Fig. 7b,c), thus suggesting that IGF2 produced by the endocrine and exocrine pancreas does not play major autocrine or paracrine growth roles that alter pancreas size. Consistent with this hypothesis, single *Igf2*^+/fl^; *Ptf1a*^Cre/+^ knockouts have normal pancreas sizes and body weights (Fig. 7b,c).

Glucose homeostasis, assessed by oral glucose tolerance tests (OGTT), was unaltered in single *Igf2*^+/fl^; *Nkx3.2*^Cre/+^ and *Igf2*^+/fl^; *Ptf1a*^Cre/+^ knockouts, as well as in double *Igf2*^+/fl^; *Nkx3.2*^Cre/+^; *Ptf1a*^Cre/+^ knockouts compared to *Igf2*^+/fl^ controls at 8 weeks of age (Supplementary Fig. 10a,b). Likewise, body weights and glucose tolerance were similar in beta-cell specific *Igf2*^+/fl^; *RIP*^Cre/+^ knockouts compared to *Igf2*^+/+^; *RIP*^Cre/+^ controls at 8 weeks (Supplementary Figs. 10 c,d,e). We next assessed glucose homeostasis regulation during pregnancy (representing a naturally occurring metabolic stress state) at E15 in 8 weeks old *Igf2*^+/fl^; *Nkx3.2*^Cre/+^ and *Igf2*^+/fl^; *RIP*^Cre/+^ knockout females. Pregnant *Igf2*^+/fl^; *Nkx3.2*^Cre/+^ knockout females are glucose intolerant compared to pregnant *Igf2*^+/fl^ littermate controls (Fig. 7d). However, no differences in glucose tolerance were observed between pregnant *Igf2*^+/fl^; *RIP*^Cre/+^ knockout females and pregnant *Igf2*^+/+^; *RIP*^Cre/+^ littermate controls (Fig. 7e).

## Discussion

This study uncovers novel roles for IGF signalling in the developmental regulation of adult pancreas size, with consequences for energy homeostasis and post-natal whole body-growth. We first observed that gene members of the IGF signalling pathway, which include IGF2 and IGF1 ligands, are highly enriched in the neonatal mesenchyme compared to the non-mesenchyme cells. Moreover, IGF2 is the top expressed gene in the mesenchyme, as revealed by RNA-seq profiling at Postnatal day 2 (P2). Mesenchymal cells express the highest levels of *Igf2* mRNA compared to all other pancreatic cell types from E16 to postnatal 8 weeks. At P2 we estimate, based on conditional deletions of *Igf2* from the diverse cell types, that 84-86% of total *Igf2* in the pancreas is expressed by the mesenchyme, ^~^15% by the endothelium and less than 1% is derived from the epithelium. These conditional deletion studies also show that *Igf2* is robustly imprinted (paternally expressed) in all cell types. Therefore, we hypothesized that mesenchymal IGF2/IGF signalling may play previously unanticipated roles in pancreatic development, and generated conditional specific mouse models mainly targeting levels in mesenchyme.

We found that mesenchymal IGF2 is principally required for normal growth of the exocrine pancreas. Deletion of *Igf2* from the mesenchyme cells from E9.5 leads to severe hypoplasia of the postnatal pancreas (^~^69%N at P2, ^~^67%N at P21 and ^~^79-83%N at 9 weeks of age). The acinar mass is mostly affected and is reduced, in line with the overall loss in pancreatic weight. This finding is in agreement with the reports of reduced exocrine mass in mice that lack both IGF ligands, *i.e*. mice lacking both *Igf1* and *Igf2* in all cells of the body, or their receptors, *i.e*. mice constitutively lacking both *Insr* and *Igf1r*^21^. However, the contribution of the individual ligands or receptors to the loss of exocrine mass has remained unclear, mainly because *Insr* or *Igf1r* single total knockouts result in either normal or increased exocrine mass, respectively, and single *Igf1* or *Igf2* total knockouts were not studied for pancreatic phenotypes. Our work now demonstrates that IGF2 is a key promoter factor of acinar growth during development. Crucially, we have ruled out a role for IGF2 expressed in the developing epithelium, which includes IGF2 expressed in acinar cells, as a determinant of exocrine pancreas growth. Accordingly, deletion of *Igf2* using *Ptf1a*-Cre, results in normal pancreatic growth. We therefore conclude that the effect of IGF2 on pancreas growth is restricted to either autocrine and/or paracrine roles in the mesenchyme.

We next addressed the question of how mesenchymal IGF2 regulates pancreas size. We propose that this is mainly achieved through secretion of IGF2 from the mesenchyme and signalling to the neighbouring cells types in a paracrine manner (see model in Fig. 8). This hypothesis is based on the following findings: first, IGF2 does not act in an autocrine manner in the mesenchyme, since its deletion does not lead to a loss of mesenchymal mass. In addition, we found very few transcriptomic changes in the *Igf2*-deficient mesenchyme. The absence of an autocrine role is further supported by the genetic manipulation of IGF type I receptor, the main receptor mediating cell proliferation effects of IGF ligands. We show that a mesenchyme specific deletion of *Igf1r* does not affect pancreas growth. We therefore conclude that the effect of *Igf2* is not caused by major morphological or functional changes to the mesenchyme that could indirectly affect the growth and development of neighbouring cells. Second, IGF2 regulates acinar growth in a paracrine manner. We used primary cell cultures to show that mesenchyme secretes IGF2 into the culture media, and that exogenous IGF2 promotes proliferation of acinar cells and increased enzymatic output through AKT signalling. *In vivo*, increased IGF2 activity specifically in the mesenchyme led to significant hyperplasia of the exocrine pancreas, as shown in two distinct genetic models (*H19DMD*^fl/+^; *Nkx3.2*^+/Cre^, and *Igf2r*^fl/+^; *Nkx3.2*^+/Cre^). Importantly, deletion of *Igf2* from the mesenchyme leads to a reduction in beta-cell mass (^~^82% N at P2), showing that the paracrine effects are not exclusive to the exocrine pancreas. Future studies will be necessary to identify the receptors that mediate the IGF2 actions in the acinar and beta cells.

**Figure 8:**
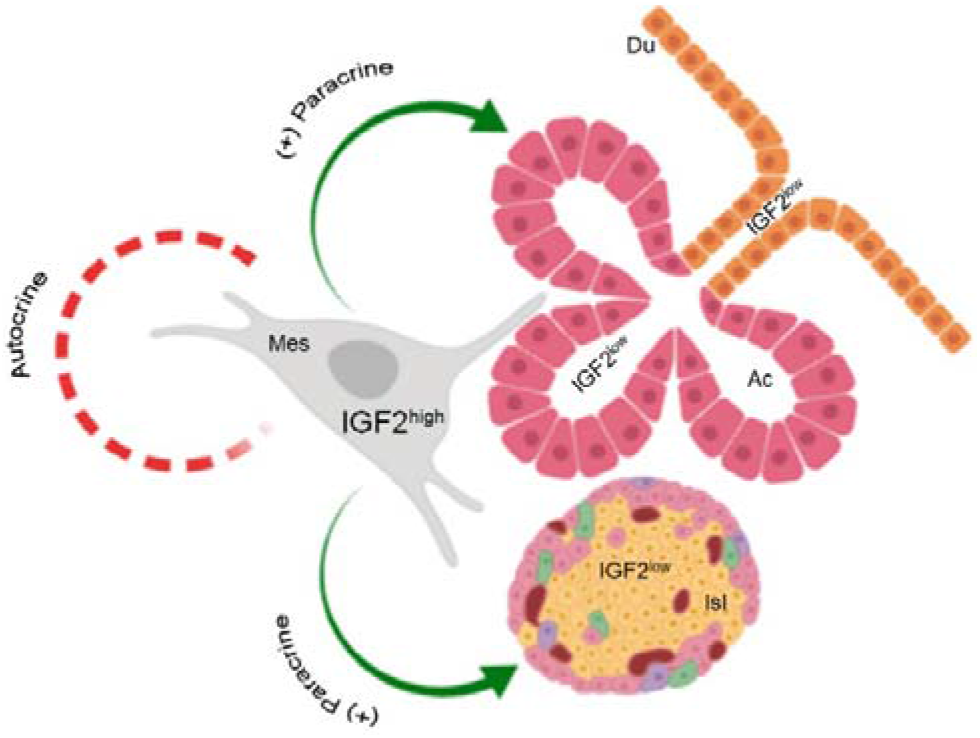
Suggested model of mesenchymal IGF2 actions in the developing pancreas. IGF2 expression is high (IGF2^high^) in the mesenchymal cells (Mes) of the pancreas and low (IGF2^low^) in other pancreatic cell types (Ac – acinar cells, Isl – pancreatic islet cells, Du – pancreatic duct cells). IGF2 produced by the mesenchymal cells exerts paracrine effects (green arrows) on pancreatic acinar cells and pancreatic islets. However, we found little evidence (red dotted line) for an autocrine role of mesenchymal IGF2 (see text). Therefore, we propose that the main function of IGF2 in the developing pancreas resides within the mesenchyme as a growth signal to the neighbouring cell types.

The lack of mesenchymal IGF2 has important physiological consequences. It is well established that the pancreas size must match the physiological demands of the host organism to promote nutrient digestion and absorption in the gut and to maintain glucose homeostasis. In this study, we report that mesenchyme *Igf2*-deficient mice have a disproportionately smaller pancreas, with a 30% size reduction at P2, which is associated with a reduced output of secreted acinar enzymes, such as lipase. Our data suggests that nutrient malabsorption due to loss of exocrine mass and/or exocrine dysfunction might be causative of the postnatal growth restriction, which is first observed in these mice at the weaning stage and maintained throughout adulthood. Mouse models with perturbations of lipase genes such as colipase (*Clps*) and pancreatic-lipase related protein 2 (*Pnliprp2*) exhibit a similar body weight growth defect phenotype due to an inability to process fat from mother’s milk^25,26^. It has been suggested that the lipase deficiency can programme a “set-point” of body weight, with an inability to catch-up in body weight later on, due to the effects of poor weight gain during the suckling period^25,26^. Interestingly, RNA-seq shows that mesenchyme *Igf2*-deficient pancreases upregulate transcription of exocrine genes involved in the production of digestive enzymes. We suggest that these changes represent adaptive responses of the small exocrine pancreas to meet the demand for digestion and absorption of nutrients in early postnatal life. However, this compensatory mechanism might not be sufficient to match demand, and whole-body growth restriction ensues. The RNA-seq studies also showed increased expression of chemokines, which are normally up-regulated by pro-inflammatory stimuli and attract immune cells by inducing a response from the innate and adaptive immune system^27,28^. Whether a pre-pancreatitis state exists, linked to the premature activation of digestive enzymes in the interstitial space of the pancreas, and how such a state might contribute to putative nutrient absorption abnormalities require further investigation. We also observed that the loss of IGF2 paracrine effects from the mesenchyme results in a modest reduction of the beta-cell mass that does not impact on glucose homeostasis control in young mice under a normal diet and physiological state. However, induced metabolic stress, as it occurs naturally during pregnancy, leads to glucose intolerance. It remains to be established if pregnancy-related beta-cell mass expansion is impaired and the extent to which other metabolic stress states, such as aging, high fat diet, acute insulin resistance states, may also uncover mesenchymal IGF2 effects on beta-cell function. Finally, we found little evidence for an autocrine growth action of IGF2 in the beta-cell that would have an impact on glucose homeostasis regulation under normal physiology. Beta-cell mass is unaffected by beta-cell specific *Igf2* deletion at 42 weeks in chow fed animals (Hammerle *et al.*, unpublished). These results are largely similar to those recently reported in 24-26 weeks old beta-cell specific *Igf2* knockout mice fed with normal chow diet^18^.

Our findings offer novel insights into mesenchymal factors that regulate pancreas organogenesis. Wnt, BMP, FGF and Hedgehog signalling pathways have been implicated as early mesenchymal factors regulating pancreas development and growth^2,29,30,31,32^, but for the majority of these studies the specific roles in the mesenchyme could not be assessed (only recently it has been possible to start interrogating the roles that genes play in the mesenchyme with the generation of specific Cre recombinases). To our knowledge, this study, using a combination of loss and gain-of-function conditional mouse models, is the first to identify a mesenchymal-specific paracrine growth signal of the developing pancreas. It also reveals that IGF signalling plays previously unappreciated roles in the mesenchyme function and in the epithelial-mesenchymal interactions during pancreas organogenesis. *Igf1*, like *Igf2*, is highly enriched in the mesenchyme compared to non-mesenchyme cells and paracrine release of IGF1 from fibroblasts, which derive from the primitive mesenchyme, stimulate acinar cell proliferation during regeneration from acute pancreatitis^33^. Interestingly, deletion of *Igf1* from the developing epithelium also results in normal exocrine pancreas growth^34^, similarly to the results reported here for *Igf2*. Further conditional knockout studies will be required to examine mechanistic aspects of the role of mesenchymal IGF1, and to establish which receptors are important as signalling transducers for IGF1 and IGF2. Our study has also highlighted a number of growth-related imprinted genes (including *Plagl1/Zac1* a master regulator of an imprinting growth network^35^) that are highly enriched in the mesenchyme, and raises the interesting hypothesis that genomic imprinting contributes significantly to mesenchymal-specific functions during pancreas development.

In summary, we report in this study that IGF2 growth actions in the pancreas are limited to the highly expressing mesenchyme, and that size-determining mechanisms are programmed in early life by the activity of this important growth factor in a paracrine fashion, with consequences for postnatal physiology. In general, this work provides novel insights into how growth factors control pancreatic architecture and is highly relevant to pancreas pathologies, from diabetes to cancer.

## Methods

### Generation of the *Igf2*^fl/+^mouse

The *Igf2* gene targeting vector carried a loxP site inserted 5’ of exon 4 and a loxP-flanked neomycin resistance cassette (neoR) inserted 3’ of exon 6 (Supplementary Fig. 1a). Details of the cloning procedures are available upon request. In brief, we used a 3.1-kb *Sal*I-*Pac*I genomic fragment (from exon 2 to intron 3 of *Igf2*) as the 5’ region of homology (5’-ROH), a 5.8-kb *Pac*I-*Afe*I genomic fragment that includes the entire *Igf2* coding sequence (exons 4 to 6) as internal ROH (Int-ROH), and a 3.1-kb *Afe*I-*Blp*I genomic fragment (3’ of exon 6) as 3’-ROH. The Int-ROH was flanked by a 5’ loxP site and a 3’ loxP-neoR-loxP sequence (Supplementary Fig. 1a). The targeting vector was linearized at a unique *Sca*I site outside the area of homology and 50 μg linearized vector were electroporated into passage 9, E14 129ola male ES cells, at 250V and 950 μF. Transfected cells were plated onto 10 gelatinized 100-mm dishes pre-seeded with fibroblast feeder cells. After 24 h in nonselective medium, cells were incubated for 8 days with G418 medium (200 μg/μl) to select for neomycin resistance. Resistant clones were picked at day 9 and expanded into 96-well plates pre-seeded with fibroblast feeder cells. We initially screened 384 G418-resistant clones by Southern blotting analysis of genomic DNA (gDNA) digested with *EcoR*I and hybridized the blots with a unique 511 bp 5’ probe (located external to 5’-ROH and obtained by PCR amplification using primers 5’Pr-F: 5’-AACAACGCGGTGGTAGGGAA-3’ and 5’Pr-R: 5’-TCAGCAGAAAAAGAAGCAGGGC-3’). Two correctly targeted clones at the 5’ end were then verified by Southern blot (*EcoR*I digested DNA) using a 590 bp 3’ probe (located external to 3’-ROH and obtained by PCR amplification using primers 3’Pr-F: 5’-ACAAAGCCCAAGACAACTCC-3’ and 3’Pr-R: 5’-CTTCCACAGTTCAAGCAACC-3’) and an additional check for multiple integrations elsewhere in the genome using a 583 bp Internal probe (located in *Igf2* exon 6 and obtained by PCR amplification with primers Int-F: 5’-AGAACCCAAGAAGAAAGGAAG-3’ and Int-R: 5’-AGAAAGACAGAACTAGCAGCC-3’). One clone with a single integration site and correctly targeted 5’ and 3’ loxP sites was thus identified (Supplementary Fig. 1b), with the loxP sequences further verified by Sanger sequencing (data not shown). The neoR cassette was excised by transiently transfecting this ES cell clone with Cre recombinase (pMC-Cre vector), followed by two rounds of subcloning. Correctly excised clones that carry a single 3’ loxP site were verified by PCR screening (480 ES subclones) and gDNA digestion with *Sph*I restriction enzyme, followed by Southern blot analysis using the same 583 bp Internal probe described above (Supplementary Figs. 1c-f). Primers used for PCR screening of *in vitro* neoR deletion were: F0 5’-TGACCTCAGCAATTCAAGTCC-3’; F1 5’-GGTAGTGGTCTTTGGCATCC-3’ and R1 5’-CAATAACTGGGGAAAAGGAGC-3’. Two independent ES clones were then microinjected into C57BL/6J blastocysts and transferred into (C57BL/6J X CBA/Ca)F1 pseudo-pregnant females to generate chimeric mice. 27 chimeras were born, all males. Germline transmitting mice were backcrossed into the C57BL/6J genetic background for more than 10 generations before being used as experimental animals. Paternal transmission of the *Igf2* floxed allele did not have any impact on fetal and placental growth kinetics (Supplementary Fig. 1g,h). Homozygous *Igf2*^fl/fl^ mice were obtained by breeding heterozygous *Igf2*^fl/+^ parents.

### Mouse strains

Targeted ES cells for the *miR-483* knock-out^22^ were generated and provided by Dr. Haydn Prosser (Sanger Institute and International Knockout Mouse Consortium Project). Details on the generation of the *miR-483* knock-out mice (depicted in Supplementary Fig. 7; Sekita *et al.*, manuscript in preparation) are available upon request. *Rosa26YFP*-stop^fl/fl^ mice^36^ were kindly provided by Dr. Martin Turner (The Babraham Institute, Cambridge). *H19DMCf*^fl/fl^ mice^23^ and *Igf2r*^fl/fl^ mice^37^ were generously provided by Prof. Bass Hassan (University of Oxford). *Igf1r*^fl/fl^ mice^38^ were imported from the Jackson Laboratory (Maine, USA). *CMV*-Cre mice^39^ were obtained from the Babraham Institute, Cambridge. This Cre recombinase is expressed soon after fertilization and allows ubiquitous deletion of floxed alleles in all tissues, including the germline^39^. *Nkx3.2*-knock-in-Cre mice^40^ were kindly provided by Dr. Warren Zimmer (Texas A&M University). *Nkx3.2* expression occurs in the mesenchyme of the developing pancreas, stomach and gut, as well as in the forming somites, but not in the endoderm-derived cells of these organs. *Nkx3.2* is expressed in the pancreatic mesenchyme as early as E9.5 and, by E12.5, its expression becomes restricted to the mesenchymal area, which will give rise to the splenic bud^40^. The *Nkx3.2*-knock-in-Cre, in which a Cre recombinase cDNA cassette and a PGK-NeoR cassette were inserted in frame within exon 1 of *Nkx3.2* (*Bapx1*) gene, faithfully replicates endogenous *Nkx3.2* expression and directs Cre activity to the foregut mesenchyme and skeletal somites starting at E9.5^2,40^. We verified the pancreatic cell-type specificity of *Nkx3.2*-Cre expression using the *Rosa26YFP*-stop^fl/fl^ reporter mice and demonstrated absence of recombinase activity in acinar cells, pancreatic endocrine cells (Fig. 1c), endothelial cells (Supplementary Fig. 4a) or duct cells (Supplementary Fig. 4b) at postnatal day P2. *Tek*-Cre transgenic mice, in which Cre expression is controlled by the endothelial-specific receptor tyrosine kinase *Tek* (*Tie2*) promoter/enhancer and therefore found uniformly in endothelial cells during embryogenesis (E7.5 onwards) and adulthood^41^, were imported from the Jackson Laboratory (Maine, USA). *Ptf1a*-knock-in-Cre mice, in which the protein-coding region of *Ptf1a* (exon 1 and 2) was replaced with a Cre-cassette, express Cre in pancreatic ducts, exocrine and endocrine cells as early as E10.5^42^. In newborn pups *Ptf1a*-Cre expression marks all acinar cells and roughly 95% of ductal and insulin-producing cells, as well as 75% of glucagon-producing cells^42^. *RIP-Cre* mice, carry a Cre transgene under the control of the rat *Ins2* (insulin 2) promoter (RIP) that directs expression to insulin-positive beta-cells from approximately E8.5-9 onwards^43^. *Ptf1a*-Cre and *RIP*-Cre strains were obtained from Central Biomedical Services (CBS Transgenic Services, University of Cambridge). All lines were bred into an inbred C57BL/6J line for >10 generations or placed into a uniform C57BL/6J genetic background using the advanced congenics program MaxBax® (The Jackson Laboratory, Maine, USA), until >98% C57BL/6J genetic contribution was achieved.

### Mouse crosses and genotyping

This research has been regulated under the Animals (Scientific Procedures) Act 1986 Amendment Regulations 2012 following ethical review by the University of Cambridge Animal Welfare and Ethical Review Body (AWERB). Mice were fed standard chow diet with 9% of kcal from fat (SDS, Essex, UK)and housed with a 12-h light/dark cycle in a temperature-controlled room (22°C). For timed mating, the day of detection of a vaginal plug was noted as embryonic day 1 (E1) and the day of birth was noted as post-natal day 0 (P0). Mice were weaned at 3 weeks of age and ear notches were used for visual identification and to collect tissue for PCR genotyping.

Mouse crosses used for each experiment are listed in Supplementary Table 1. Throughout the paper, *Igf2*^+/fl^ represents genotype of offspring that inherited the floxed allele from father; *Igf2*^fl/+^ represents genotype of offspring that inherited the floxed allele from mother; *Igf2*^+/+^ represents genotype of offspring with wild-type alleles at the *Igf2* locus; *Nkx3.2*-Cre represents genotype of offspring that are heterozygous Cre-carriers (^Cre/+^ or ^+/Cre^); *Igf2*^+/fl^; *Nkx3.2*^Cre/+^ represents offspring with mesenchymal *Igf2* deletion on the paternal allele, having inherited the floxed allele from father and the Cre allele from mother; *Igf2*^fl/+^; *Nkx3.2*^+/Cre^ represents offspring with mesenchymal *Igf2* deletion on the maternal allele, having inherited the floxed allele from mother and the Cre allele from father; *H19DMD*^fl/+^; *Nkx3.2*^+/Cre^ represents offspring with mesenchymal *H19DMD* deletion on the maternal allele, having inherited the floxed allele from mother and the Cre allele from father. All strains were genotyped by standard PCR using DNA extracted from ear biopsies (adult mice) or tail DNA (embryos or post-terminal). PCR was performed using the Red Taq Ready PCR system (Sigma Aldrich) using primers described in Supplementary Table 2, followed by separation of PCR amplicons by agarose gel electrophoresis.

### Southern blotting screening of ES clones

gDNA were digested with *EcoR*I or *Sph*I restriction enzymes and electrophoresed on 0.8% agarose gels in 1×TBE buffer, alkaline blotted onto Hybond N+ membranes (Amersham), and UV cross-linked (Stratalinker, Stratagene). Probes were obtained by PCR and radiolabeled (α-32P-CTP). Hybridisation and washing of Southern blots was performed as described^44^. Membranes were exposed overnight to MS film (Kodak).

### Northern blotting analysis of *Igf2* mRNA

Total RNA (10 μg) extracted from E19 placenta and liver samples using RNeasy midi kits (Qiagen) according to manufacturer’s protocol, was separated in low-percentage formaldehyde gels, blotted onto Nytran-plus membrane (Schleicher and Schuell), and UV cross-linked (Stratalinker, Stratagene). The RNA blots were hybridized with radiolabelled (α-32P-UTP) *Igf2* and *Gapdh* cDNA probes. We carried out hybridization and washing of northern blots as described^45^. Transcript levels were quantified by PhosphorImager analysis (Molecular analyst software, Biorad).

### Western blotting analyses

For Western blot analysis of IGF2, 5 μl serum samples collected at E19 were loaded into Bis-Tris gels (NuPAGE Novex, Life technologies) and transferred to nitrocellulose membranes (Invitrogen). 4 ng human recombinant IGF2 (292-G2-050, R&D Systems) was used as positive control, and the Novex Sharp protein ladder (Life technologies) was used as marker. The membranes were initially stained with Ponceau red (Invitrogen) and imaged with the Biorad GelDoc system, the intensity of bands being used as internal control for protein loading. The membranes were then blocked with 5% skimmed milk in TBS-T for 1 h at 4°C, after which were incubated overnight at 4°C with goat antihuman IGF2 antibody (R&D systems AF-292) diluted 1: 1,000 in TBS-T containing 0.3% skimmed milk. After 4×10 min washes with milliQ water, the blots were incubated for 1h at room temperature with the secondary antibody (1:2,000 rabbit anti-goat IgG coupled to HRP, SantaCruz) in TBS-T containing 3% skimmed milk. Blots were washed 4×10 min with milliQ water, exposed to substrate (Clarity ECL Western Blotting Substrate, Biorad) for 5 minutes and imaged with the Biorad GelDoc system. Bands were quantified using the ImageLab software (Biorad), the output values being normalized to the corresponding Ponceau red loading control values.

For Western blot analysis of AKT phosphorylation, acinar cell pellets were lysed in 30μl RIPA containing protease and phosphatase inhibitors. Protein concentration was determined by DC assay (Biorad) and 10μg/lane total cell lysates were electrophoresed through 4-12% Bis-Tris NuPAGE gels with MOPS running buffer. Proteins were transferred to nitrocellulose by iBlotII (P3 8:00 min) prior to being blocked for an hour in 3%BSA/TBST. Primary antibody to pAKT (S473) [Cell Signalling Technology (CST) 9271], total AKT (CST 2920), or Beta-actin (CST 4967) was incubated overnight at 4°C. Membrane was washed 5 times before detection of bound antibody by detection of chemiluminescent signal associated with HRP-conjugated anti-rabbit or anti-mouse secondary antibody. Membrane was washed 5 times before detection of bound antibody by detection of chemiluminescent signal associated with HRP-conjugated secondary antibody.

### Pancreas immunostainings, imaging and stereology analysis

Pancreata were dissected under microscope (for embryonic and early post-natal analyses), fixed in 4% paraformaldehyde in PBS overnight, dehydrated and then embedded in paraffin. Paraffin blocks were cut at 5 μm thickness, sections were then deparaffinised, rehydrated, stained and mounted with coverslips. Conditions used for antigen retrieval, blocking and combinations of primary and secondary antibodies used for each staining are described in Supplementary Table 3. Counterstaining was performed with haematoxylin for light microscopy stains or DAPI for fluorescent stains.

Light microscopy stained slides were imaged using the NanoZoomer whole slide scanner (Hamamatsu). Fluorescent immunostainings were imaged with a LSM510 Meta confocal laser scanning microscope and the ZEN 2009 software (Carl Zeiss, Germany) or scanned using an Axio whole slide scanner. Whole slide scans of stained sections (seven sections spaced at 100 μm distance for each P2 pancreas) were analysed using the Visiopharm automated image quantification software (Visiopharm, Denmark) to measure the area of positive cell-specific staining or to count individual nuclei. The mass of each cell-type was calculated using the formula:

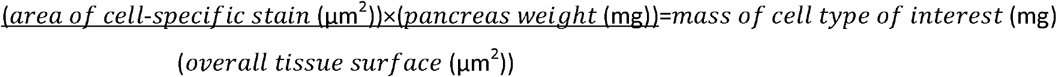

### mRNA in situ hybridization (ISH) for *Igf2*

ISH was performed as described^46^, with minor modifications. Briefly, a region of 415 bp spanning coding *Igf2* exons 4-6 was amplified by PCR using primers: F: 5’-CACGCTTCAGTTTGTCTGTTCG-3’ and R: 5’-GCTGGACATCTCCGAAGAGG-3’ and cDNA from whole P2 pancreata as template. The PCR product was cloned into a pCR2.1-TOPO plasmid (Invitrogen) containing M13 primers and a T7 RNA polymerase transcription initiation site. Sense (S) and antisense (AS) RNA probes were generated and labelled with Digoxigenin (DIG) by *in vitro* reverse transcription, according to manufacturer’s instructions (Roche). Pancreata were dissected in cold phosphate buffered saline (PBS) and fixed overnight in 4% paraformaldehyde in 0.1% diethylpyrocarbonate (DEPC)-PBS at 4°C. After rinsing in DEPC-PBS, tissues were dehydrated and then embedded in paraffin in RNase-free conditions. Pancreas sections (7 μm thick) mounted on polysine slides (VWR) were de-waxed, rehydrated in PBS, post-fixed in 4% paraformaldehyde for 10 minutes, treated with proteinase K (30 μg/ml) for 10 min at room temperature, acetylated for 10 minutes (acetic anhydride, 0.25%) and hybridized overnight at 65°C in a humidified chamber with DIG-labeled probes diluted in hybridization buffer (containing 200 mM sodium choride, 13 mM tris, 5 mM sodium phosphate monobasic, 5 mM sodium phosphate dibasic, 5 mM EDTA, 50% formamide, 10% dextran sulfate, 1 mg/ml yeast tRNA and 1× Denhardt’s [1% w/v bovine serum albumin, 1% w/v Ficoll, 1% w/v polyvinylpyrrolidone]). Two 65°C posthybridization washes (1× SSC, 50% formamide, 0.1% tween-20) followed by two room temperature washes in 1× MABT (150 mM sodium chloride, 100 mM maleic acid, 0.1% tween-20, pH7.5) were followed by 30 minutes RNAse treatment (400 mM sodium chloride, 10 mM tris pH7.5, 5 mM EDTA, 20 μg/ml RNAse A). Sections were blocked for 1 hour in 1×MABT, 2% blocking reagent (Roche), 20% heat inactivated goat serum and then incubated overnight with anti-DIG antibody (Roche; 1:2,500 dilution) at 4°C. After 4×20 min washes in 1× MABT, slides were rinsed in 1× NTMT (100 mM NaCl, 50 mM MgCl, 100 mM tris pH 9.5, 0.1% tween-20) and incubated with NBT/BCIP mix in NTMT buffer, according to manufacturer’s instructions (Promega). Slides were counterstained with nuclear fast red (Sigma), dehydrated and cleared in xylene and mounted in DPX mounting medium (Sigma). Pictures were taken with a camera attached to a light microscope.

### Fluorescence-activated cell sorting (FACS)

Mice were sacrificed by decapitation (embryos and P2 neonates) or cervical dislocation (P5 neonates and older). Then, pancreata were dissected and dissociated into single cells using trypsin-EDTA (Sigma Aldrich) at 37°C for 20 min. After one wash with ice-cold PBS, the cells were passed through 70 μm strainers and single-cell suspensions were sorted into YFP-positive and YFP-negative fractions using an Aria-Fusion cell sorter (BD Bioscience). Dead cells were excluded based on forward and side scatter profiles and the uptake of 7AAD (7-Aminoactinomycin D dead cell stain, Life Technologies). Sorted YFP-positive and YFP-negative cells were pelleted by centrifugation and flash frozen in liquid nitrogen, then stored at −80°C until use.

### Total RNA and microRNA extraction (pancreas and sorted cells)

Total RNA was extracted from whole pancreata, cells isolated by FACS or cultured cells using the RNEasy Mini kit (Qiagen) or the RNEasy Micro kit (Qiagen), respectively, according to manufacturer’s instructions, with additional removal of contaminating DNA using QIASpin DNA eliminator columns. microRNA was extracted using the miRNeasy kit (Qiagen), according to manufacturer’s instructions with an additional step of DNaseI treatment (Qiagen).

### Quantitative real-time PCR (qRT-PCR)

Concentration and integrity of total RNA was verified by NanoDrop (Thermo Scientific) and agarose gel electrophoresis, respectively. For RNA extracted from whole pancreata, reverse transcription was performed using the RevertAid RT Reverse Transcription Kit (Life technologies), according to manufacturer’s instructions. In the case of total RNA extracted from sorted cells, cDNA was produced using the QuantiTect Whole Transcriptome Kit (Qiagen) following manufacturer’s instructions. Primers used for qRT-PCR are listed in Supplementary Table 4. Annealing temperatures were tested by gradient PCR using pancreas-cDNA as a template. qRT-PCR was performed using the SYBR Green JumpStart Taq Ready Mix (Sigma Aldrich) on an ABI Prism 7900 system (Applied Biosystems). Gene expression levels were normalized to the housekeeping gene *Ppia* (peptidylpropyl isomerase A or cyclophilin-A), which has previously been established as a good housekeeping gene for pancreas gene expression studies^47^ and is stably expressed between various developmental time points (data not shown). For microRNAs, reverse transcription was performed using the TaqMan Micro RNA Reverse Transcription kit (Applied Byosystems), according to manufacturer’s protocol. qRT-PCR was performed using TaqMan assays (TM: 002560 for mmu-miR-483; TM: 001232 for snoRNA202 and TM: 001234 for snoRNA234) and TaqMan 2x Universal PCR Master Mix (Applied Byosystems). Levels of expression were calculated using the 2^−ΔΔCt^ method^48^.

### Complementary DNA library preparation and RNA-seq analysis

Input RNA for genome-wide transcriptome analysis was verified for concentration and quality using Agilent RNA Pico chips, according to manufacturer’s instructions. All RNA samples had RNA integrity numbers (RIN) >7.5. Total RNA (2 ng) was whole-transcriptome amplified using the Ovation RNA–seq System V2 (NuGEN). To prepare the RNA–seq libraries, the amplified cDNA (2 μg per sample) was fragmented to 200 bp using a Bioruptor Sonicator (Diagenode), and barcode ligation and end repair were performed using the Ovation Rapid DR Library System (NuGEN). The barcoded libraries were combined and loaded onto an Illumina HiSeq 2500 system for single-end 50-bp sequencing at the Genomics Core Facility, Cambridge Institute, CRUK. The reads were aligned onto the mouse GRCm38 genome using TopHat 2.0.11^49^. Gene abundance and differential expression were determined with Cufflinks 2.2.1^50^ and expressed in fragments per kilobase per million mapped reads (FPKM). The cut off for expression was set at ≥ 1 FPKM. Genes with a linear fold expression change greater than 1.5 and a Benjamini–Hochberg false discovery rate <5% were considered differentially expressed.

### Functional annotation and enrichment analysis

DAVID (Database for Annotation, Visualization and Integrated Discovery; v6.8 http://david.abcc.ncifcrf.gov/, accessed March 2017) was performed to assess whether there was enrichment for genes implicated in particular biological processes within the differential expressed gene lists identified by RNA-seq. Enriched gene ontology (GO) terms with an FDR□<□0.05 were considered significant. These terms were then clustered semantically using REViGO (Reduce and Visualize GO)^51^, which removes redundancy. The results obtained by REViGO were ordered according to log10 p values.

### Whole pancreas lipase content and amylase measurements

For whole pancreas lipase measurements, frozen samples were removed from −80°C and thawed on ice. 100 μL TK lysis buffer with proteinase inhibitors (Calbiochem) was added to each pancreas and tissues were disrupted on ice with a homogenizer (Biospec Tissue Tearor). Lysates were spun at 3,000 rpm for 15 min at 4°C and supernatants were used for analysis. Lipase levels were measured using a lipase activity assay (Dimension RXL, Siemens).

For amylase measurements, cultured acinar cells were lysed in 100 μL RIPA buffer, spun at 3,000 rpm for 15 min at 4°C and the supernatants used for analysis. Amylase levels were measured in an autoanalyser (Siemens Dimension RXL) through a colorimetric reaction based on the ability of amylase to hydrolyse the chromogenic substrate 2-chloro-4-nitrophenol linked with maltotriose (CNPG3) into the coloured product 2-chloro-4-nitrophenol (CNP) plus CNPG2, maltotriose G3 and glucose. The absorbance increase at 405 nm is proportional to the amylase activity in the sample. Amylase levels were normalized to the total protein content determined by a BCA assay (Pierce BCA Protein Assay Kit, Thermo Fisher Scientific, 23225).

### Primary pancreatic acinar and mesenchymal cell isolation and culture

Primary pancreatic cells were isolated as previously described^52^ and adapted here to P2 samples. Briefly, pancreata from an entire P2 litter were micro-dissected under microscope and sterile conditions and pooled in one tube containing HBSS, 0.25 mg/ml of trypsin inhibitor and 1% Penicillin-Streptomycin mix. After rinsing, pancreata were digested for 20-30 min at 37°C with a collagenase IA solution (HBSS containing 10mM HEPES, 200 U/ml of collagenase IA, and 0.25 mg/ml trypsin inhibitor). The digestion was stopped by placing the samples on ice and addition of FBS (fetal bovine serum) for a 2.5% final concentration. After additional washing steps, the cell suspension was passed through a 100 μm cell strainer, allowing the passage of acini and mesenchymal cells, while retaining the non-digested fragments and larger pancreatic islets. The cells were placed in the culture media (basal Waymouth’s media containing 2.5% FBS, 1% Penicillin-Streptomycin mix, 0.25 mg/ml trypsin inhibitor, and 25 ng/ml of recombinant human EGF) and plated into six-well plates pre-coated with poly-L-lysine. After one hour incubation at 37°C under 5% (v/v) CO2 atmosphere, the floating acini were transferred to a new well, while the mesenchyme cells remained attached. The cells were cultured under the above conditions for up to four days, with daily media changes.

### Measurement of IGF2 secretion by primary mesenchymal and acinar cells *in vitro*

The primary mesenchymal and acinar cells were isolated and cultured as described above until reaching confluency. Media (1mL) was collected 24h post-confluency and freeze-dried using a Christ Gamma 2-16 LSC Freeze dryer. Dry pellets were re-dissolved in 50 μl RIPA buffer and IGF2 measured by ELISA (Mouse IGF-II DuoSet ELISA kit, R&D Systems – DY792) using an assay adapted for the MesoScale Discovery electrochemiluminescence immunoassay platform (MSD), as recently described^53^.

### *In vitro* IGF2 treatments

To assess the impact of exogenous IGF2 on intracellular signalling via AKT, freshly isolated acinar cells were starved for two hours in basal Waymouth’s media, without FBS and EGF. IGF2 treatments (with vehicle or 1 nM, 10 nM or 50 nM mouse recombinant IGF2) were performed for 10 min at 37°C under 5% (v/v) CO2 atmosphere, then the cells were immediately placed on ice, washed two times in ice-cold PBS, pelleted and flash-frozen on dry-ice and stored at −80°C until analysed by Western blotting.

To assess the impact of exogenous IGF2 on amylase production by acinar cells, subconfluent acinar cultures (48 hours after isolation) were placed in basal Waymouth’s media containing 1% Penicillin-Streptomycin mix and 0.25 mg/ml of trypsin inhibitor. FBS was replaced with Serum Replacement (Sigma Aldrich S0638) that does not contain any IGF2. The treated cells received 50 ng/ml mouse recombinant IGF2 for 48 hours, with new media added every day. After the treatment, cells were harvested and dissociated into single-cell suspension using trypsin, counted with a Cedex XS Analyser (Roche), washed, pelleted and stored at −80°C until used for amylase measurement.

### Oral glucose tolerance test

Conscious mice were used for an oral glucose tolerance test after 6 hours of fasting (from 8am to 2pm). Blood samples (≤5 μL) were taken from the tail vein immediately before administration of glucose by oral gavage (20% weight for volume, 2g/kg body weight based on average body weights for each experimental group) and thereafter at the time points indicated, and used for glucose measurements with a glucose meter (AlphaTRAK).

### Statistical analysis

Statistical analysis was performed using GraphPad Prism 7 software. For two groups with up to 6 samples, statistical analysis was performed using Mann-Whitney or the Wilcoxon matched-pairs signed rank test. For two groups with ≥6 samples, to determine whether the data was of a Gaussian distribution, the Shapiro-Wilk test was first applied, followed by un-paired or paired Student’s t-tests, as appropriate. Where more than two groups exposed to the same treatment were analysed, 1-way ANOVA with Dunnett’s post-hoc or Friedman’s test with Dunn’s correction tests were used, comparing every mean to a control mean. OGTT data was analysed using two-way ANOVA with Sidak’s multiple comparison tests, using the time from administration and the genotype as factors (two genotypes) or by repeated measures 1-way ANOVA with Dunnett’s post hoc test for multiple comparisons against the controls (four genotypes). The area under the curve (AUC) was calculated by the trapezoidal rule and used for analyses by unpaired Student’s t tests (two genotypes) or by 1-way ANOVA with Dunnett’s post-hoc tests (four genotypes). Unless stated otherwise, data is-shown as average values and error bars represent SEM or SD. N.S. p >0.05; * p ≤0.05; ** p ≤0.01, *** p ≤0.001.

### Data availability

RNA-seq data have been deposited in the Gene Expression Omnibus (GEO) under the accession number GSE100981. Other data and materials are available upon request from the corresponding author.

## Acknowledgements

This work was supported by Biotechnology and Biological Sciences Research Council (grant BB/H003312/1 to M.C., S.E.O., A.V.P.); the Medical Research Council ([MRC_MC_UU_12012/4 and MRC_MC_UU_00014/4] to M.C and S.E.O, [MRC 979241] to C.H., [MRC_MC_UU_12012/5] to Metabolic Diseases Unit), the Wellcome Trust ([Strategic Award 100574/Z/12/Z], [102355/Z/13/Z] to N.M.S.) and [098051] for *miR-483* knockout production), the NIHR Cambridge BRC Cell Phenotyping Hub (in particular we wish to thank Natalia Savinykh for help with flow cytometry cell sorting) and the Spanish Ministry of Economy and Competitiveness (grants BFU2012-33594 and BFU2013-47384-R to G. M.-G.). We thank Debbie Drage, Martin George and in particular Ted Saunders (The Babraham Institute Gene Targeting Facility) for help with generating the *Igf2*^+/fl^ mice; Adrian Wayman (West Forvie Phenomics Center) and Adriana Izquierdo-Lahuerta and Yurena Vivas (Universidad Rey Juan Carlos) for help with mouse husbandry; Keli Philips and Gregory Strachan (Imaging Core facility), for help with preparing tissue samples for histology and confocal microscopy imaging, respectively; Keith Burling (Biochemical Assay Laboratory) for performing lipase, amylase and IGF2 measurements; Dan Hart (MRL Disease Model Core) for performing oral glucose tolerance tests; Dr. Claire Stocker from the University of Buckingham for providing training on pancreas stereology; Dr. Allan Bradley for providing *miR-483* knockout ES cells.

## Contributions

C.M.H. performed most of the experiments, contributed to the design of the study and wrote the manuscript. I.S. generated the conditional *Igf2* knock-out mouse, contributed experimentally to most of the experiments and their design and wrote the manuscript. G.V.B. contributed to the *in vitro* experiments on pancreatic acinar cells. N.M.S. contributed to the analysis of the mouse phenotypes (in particular imaging, stereological measurements and OGTT analyses during pregnancy). C.M.H., I.S., G.V.B., N.M.S., B.Y.H.L., W.N.C, G.M.-G. and M.C. analysed the data. W.E.Z. generated and provided the *Nkx3.2-Cre* mouse model and I.Z., H.M.P. and Y.S. contributed to the *miR-483* mouse model study. M.M. prepared RNA-seq libraries. A.V.-P., S.E.O., G.M.-G., and M.C. discussed and initiated the study. M.C. designed and led the study, and wrote the manuscript, with contributions to the final version from all authors.

## Competing interests

The authors declare no competing financial interests.

---

**Supplementary Figure 1:**
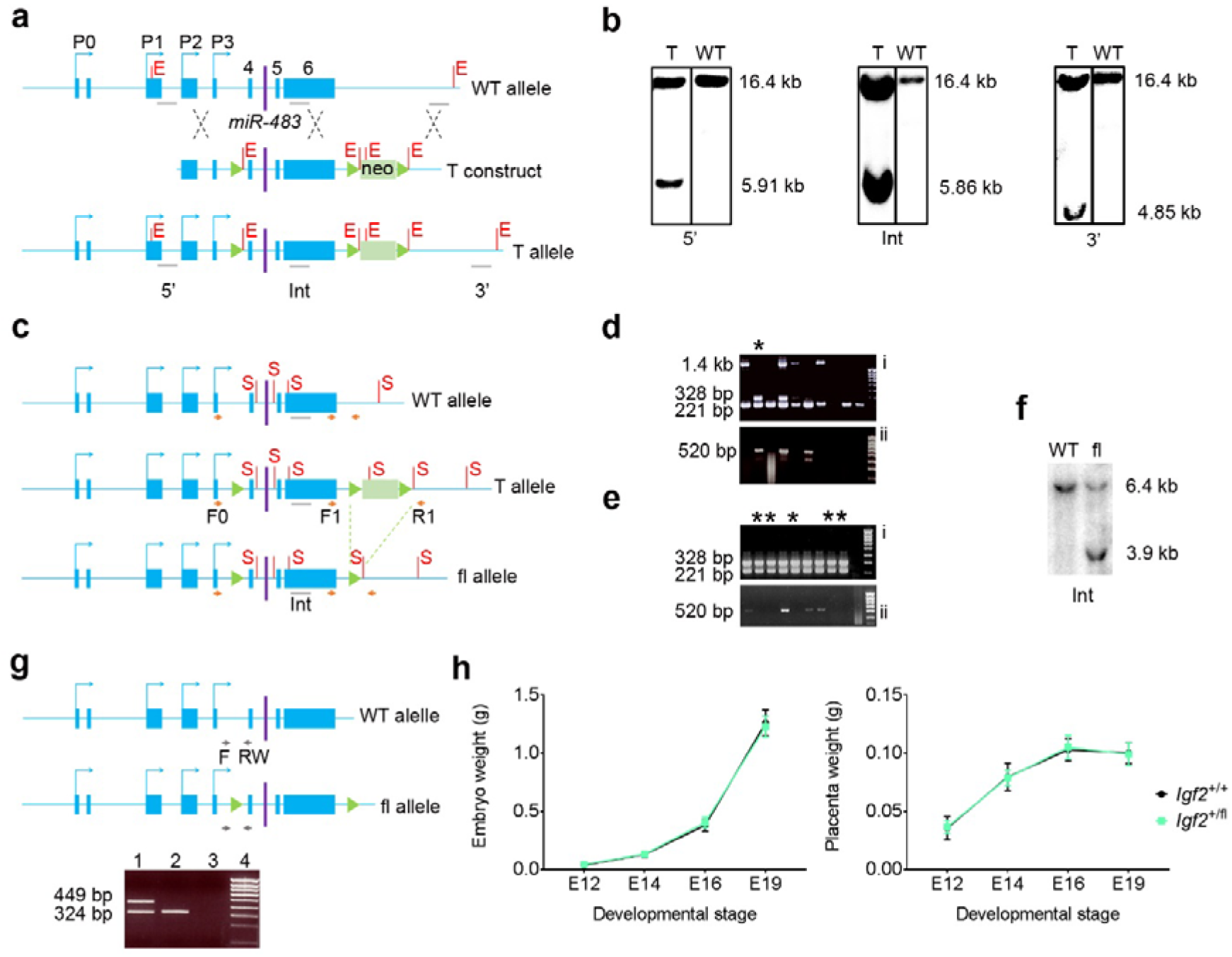
Generation of *Igf2* conditional knockout mice. (see Methods for further details). **(a)** Targeting strategy to generate an *Igf2* allele with coding exons 4 to 6 flanked by loxP sites (not drawn to scale). Blue boxes – exons; P0-P3 – alternative promoters; green triangles – loxP sites; neo – neomycin cassette; WT – wild-type; T – targeting (construct); E – EcoRI restriction sites; 5’, Int, 3’: location of 5’, internal and 3’ Southern blotting probes, respectively **(b)** Southern blot confirmation of homologous recombination between targeting vector and endogenous *Igf2* sequences in genomic ES cell DNA, digested with EcoRI and hybridized with 5’, Int or 3’ probes. Diagnostic molecular weights (kb) are indicated in each panel. T and WT – targeted and wild-type clones, respectively. **(c)** Screening strategy for loxP recombination events (not drawn to scale). Correctly targeted ES clones (as shown in (b)) that are transiently exposed to Cre recombinase *in vitro* will undergo three possible independent recombination events involving the loxP sites, which can be discriminated by PCR and Southern blotting. F0, F1, R1 – PCR screening primers; S – SphI restriction sites; Int – Internal Southern probe (d) Five 96-well plates containing targeted ES cells transfected with Cre recombinase were screened by F1+R1 primer PCR (panel i) or F0+R1 primer PCR (panel ii). F1+R1 PCR products of 221 bp, 1.4 kb and 328 bp are diagnostic of the wild-type allele, neomycin cassette, and neomycin cassette deletion, respectively (panel i). F0+R1 PCR products of 520 bp are diagnostic of a deletion that includes the neomycin cassette and exon4-6 region (in panel (ii)). Since all clones that had deletion of the neomycin cassette (*i.e*. 328 bp) also had cells with deletion of exon4-6 region (520 bp), the clone indicated with a star in panel (i) was subsequently subcloned for selection of cells with neomycin cassette excision events only. **(e)** Representative ES subclones (starred) with deletion of the neomycin cassette only (*i.e*. 328 bp in panel (i) but absence of 520 bp PCR product in panel ii). **(f)** Southern blot confirmation of deletion of the neomycin cassette in ES subclones identified in **(e)**. DNA was digested with SphI and hybridized with the internal probe: 6.4 kb – wild-type allele (WT); 3.9 kb – floxed allele (fl), resulting from Cre-induced deletion of the neomycin cassette. (g) PCR genotyping of *Igf2*^+/fl^ mice was performed using primers F and RW flanking the 5’ loxP site (primer sequences are shown in Supplementary Table 2). A representative example of tail DNA PCR is shown for mice carrying one floxed *Igf2* allele (lane 1) wild-type littermate (lane 2); lane 3 – no template PCR control; lane 4 – 100 bp DNA ladder. (h) Carriers of an *Igf2* floxed allele that was inherited paternally show identical embryonic and placental weights as littermate controls (E12: n=8 wild-type and n=3 floxed; E14: n=16 wild-type and n=7 floxed; E16 n=12 wild-type and n=10 floxed; E19: n=29 wild-type and n=24 floxed). Data is shown as average weight; error bars represent standard deviation (SD).

**Supplementary Figure 2:**
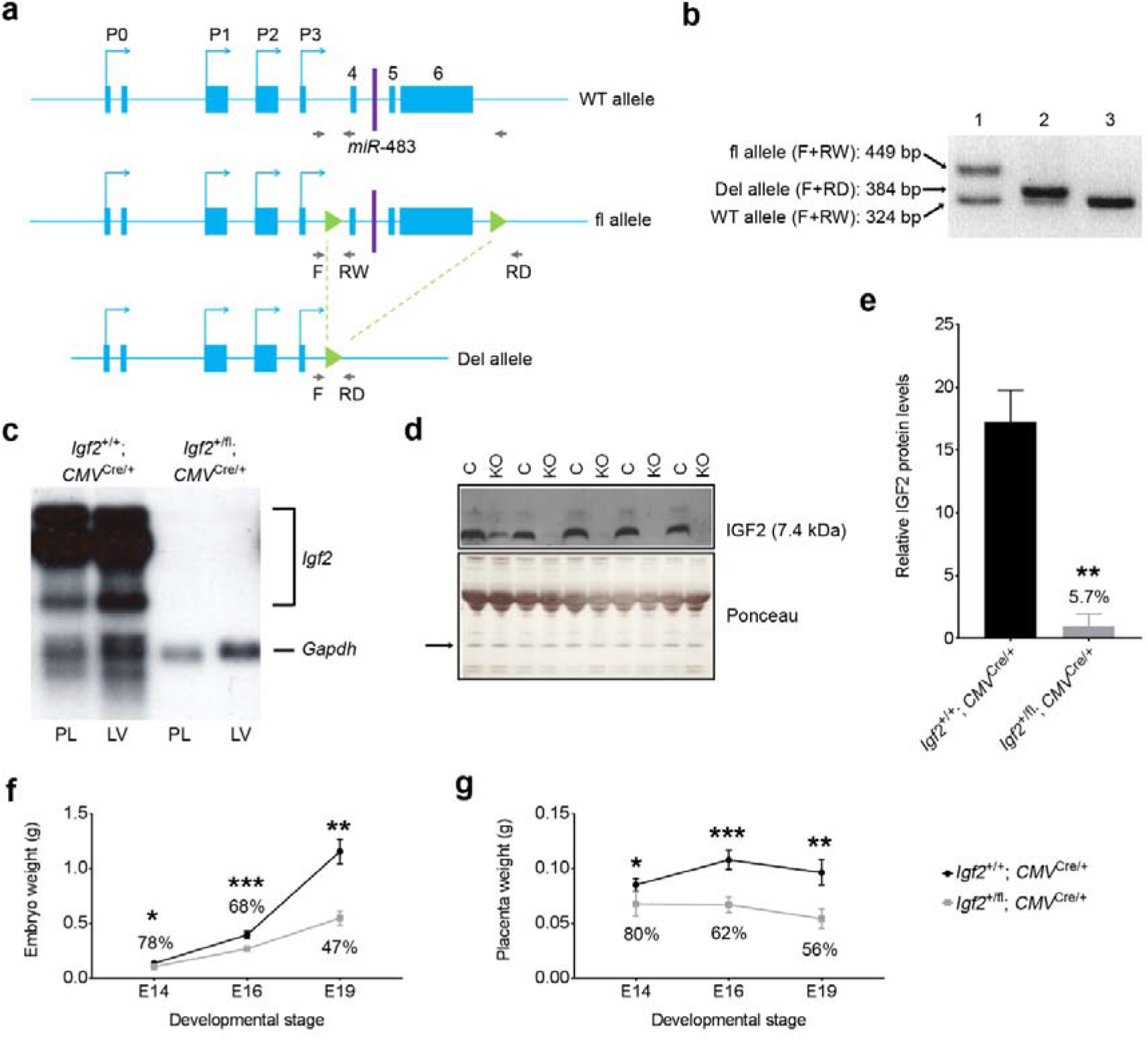
Cre-mediated deletion of the *Igf2* floxed allele. **(a)** PCR strategy to identify deletion events at DNA level. PCR products obtained in a tri-primer (F+RW+RD) PCR are diagnostic of the wild-type allele (WT: 324bp), floxed allele (fl: 449 bp) and deleted allele (Del: 384bp). Representative tail DNA PCR examples are shown in (b) for *Igf2*^+/fl^ mice that carry floxed and wildtype alleles (lane 1), *Igf2^+/fl^*; CMV^Cre/+^ mice carrying deleted and wild-type alleles (lane 2), *Igf2*^+/fl^ control littermates with wild-type alleles only. **(c)** to (g) *CMV-Cre* mediated deletion of the *Igf2* floxed allele *in vivo*. Heterozygous floxed *Igf2* males were mated with females homozygous for *CMV*-Cre (active in all cells of embryos and placentae), and offspring analysed for levels of *Igf2* deletion by northern blotting **(c)**, Western blotting **(d)** and **(e)**, and growth curves from E14 to E19 of gestation **((f)** and **(g)). (c)** Northern blot analysis of *Igf2* mRNA levels, showing wild-type levels of expression in controls *Igf2*^+/+^; *CMV*^Cre/+^) in both placenta (PL) and liver (LV) at E19, and absence of all *Igf2* transcripts upon CMV-Cre mediated deletion of the paternally inherited floxed allele (*Igf2*^+/fl^; *CMV*^Cre/+^). *Gapdh* – internal control for RNA loading. (d) Western blotting analysis of the mature form of IGF2 in serum samples collected from E19 controls (C: *Igf2*^+/+^; *CMV*^Cre/+^) and mutant (KO: *Igf2*^+/fl^; *CMV*^Cre/+^) embryos. Normalisation of IGF2 expression across samples was performed against Ponceau-stained protein band (arrow), and the relative quantification of IGF2 levels for the two genotypes is shown in **(e)**. Data is shown as average values; error bars represent SEM; ** p<0.01 by Mann-Whitney test. **(f)** and **(g)** Mice with a *CMV*-Cre mediated deletion of the paternally inherited *Igf2* floxed allele (*Igf2*^+/fl^; *CMV*^Cre/+^) show a similar embryonic **(f)** and placenta **(g)** growth phenotype to *Igf2* null mice, i.e. ~ half of the weight of littermate controls *Igf2*^+/+^; *CMV*^Cre/+^) at the end of gestation (E14: n= 3 litters; E16: n=9 litters; E19: n=4 litters). Data is shown as average values; error bars represent SD; * p<0.05; ** p<0.01; *** p<0.001 using paired student t tests.

**Supplementary Figure 3:**
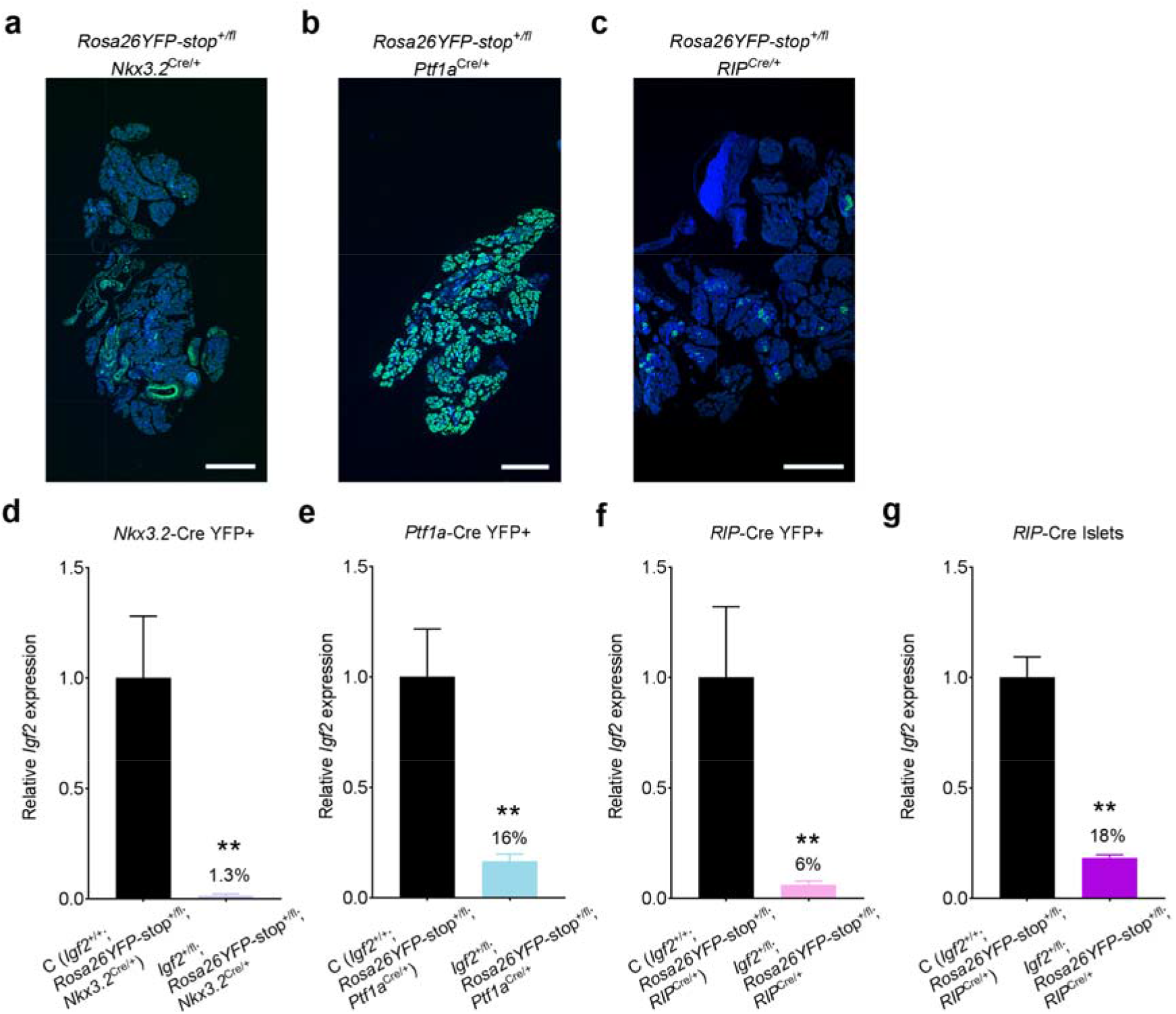
Assessment of specificity and efficiency of pancreas-expressing Cre lines using the *Rosa26YFP*-stop^fl/fl^ reporter mouse, at postnatal day 2 (P2). **(a), (b)** and **(c)** Confocal fluorescence microscopy images of sections stained for the YFP protein (green) and nuclei (DAPI, blue). Upon Cre-mediated deletion of the loxP-Stop-loxP cassette, the YFP protein is expressed in the pancreatic mesenchyme (*Nkx3.2*-Cre), pancreatic epithelial cells (*Ptf1a*-Cre) and pancreatic beta cells *(RIP-Cre)*. Scale bar: 500 μm. **(d), (e)** and **(f)** *Igf2* mRNA expression measured by qRT-PCR in YFP+ cells collected from offspring with the genotypes indicated. **(g)** *Igf2* mRNA expression measured by qRT-PCR in pancreatic islets isolated from 11-week old mice with the genotypes indicated. For panels **(d)** to **(g)** expression data was normalized to *Ppia* and shown as averages + SEM relative to levels measured in control (C) littermates, arbitrarily set to 1. Percentage values indicate the level of *Igf2* mRNA reduction relative to controls (n=4-6 samples/genotype; ** p<0.01 by Mann-Whitney tests).

**Supplementary Figure 4:**
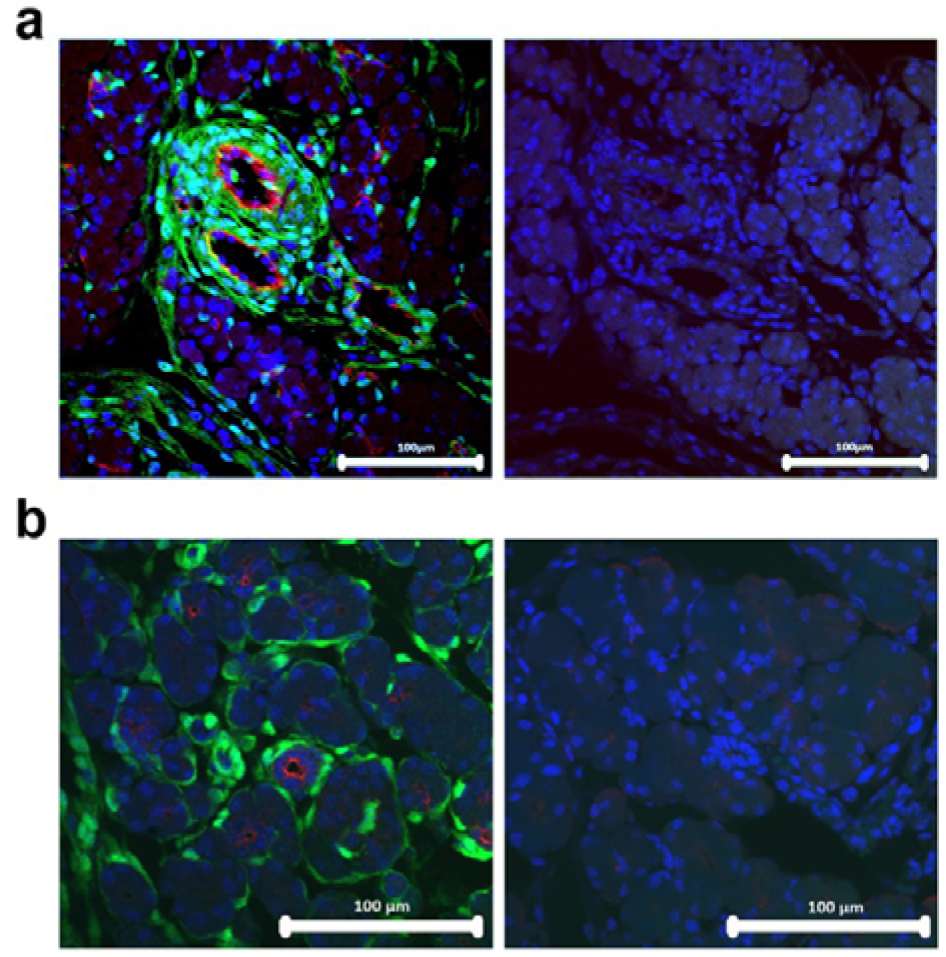
Assessment of mesenchyme-specific activity of *Nkx3.2*-Cre at postnatal day 2 (P2). Representative immunofluorescence for **(a)** endothelial cell marker CD31 (red) and **(b)** duct cell marker pan-cytokeratin (red) showing no stain overlap with the *Nkx3.2-Cre* driven YFP expression as a marker of mesenchyme cells (green). DAPI (blue) stains the nuclei. *Nkx3.2*-Cre is therefore not expressed in endothelial and duct cells, being confined to mesenchymal cells surrounding other cell types, as reported before by others. Panels on the right show negative controls performed in consecutive sections, without primary antibodies. Scale bars: 100 μm.

**Supplementary Figure 5:**
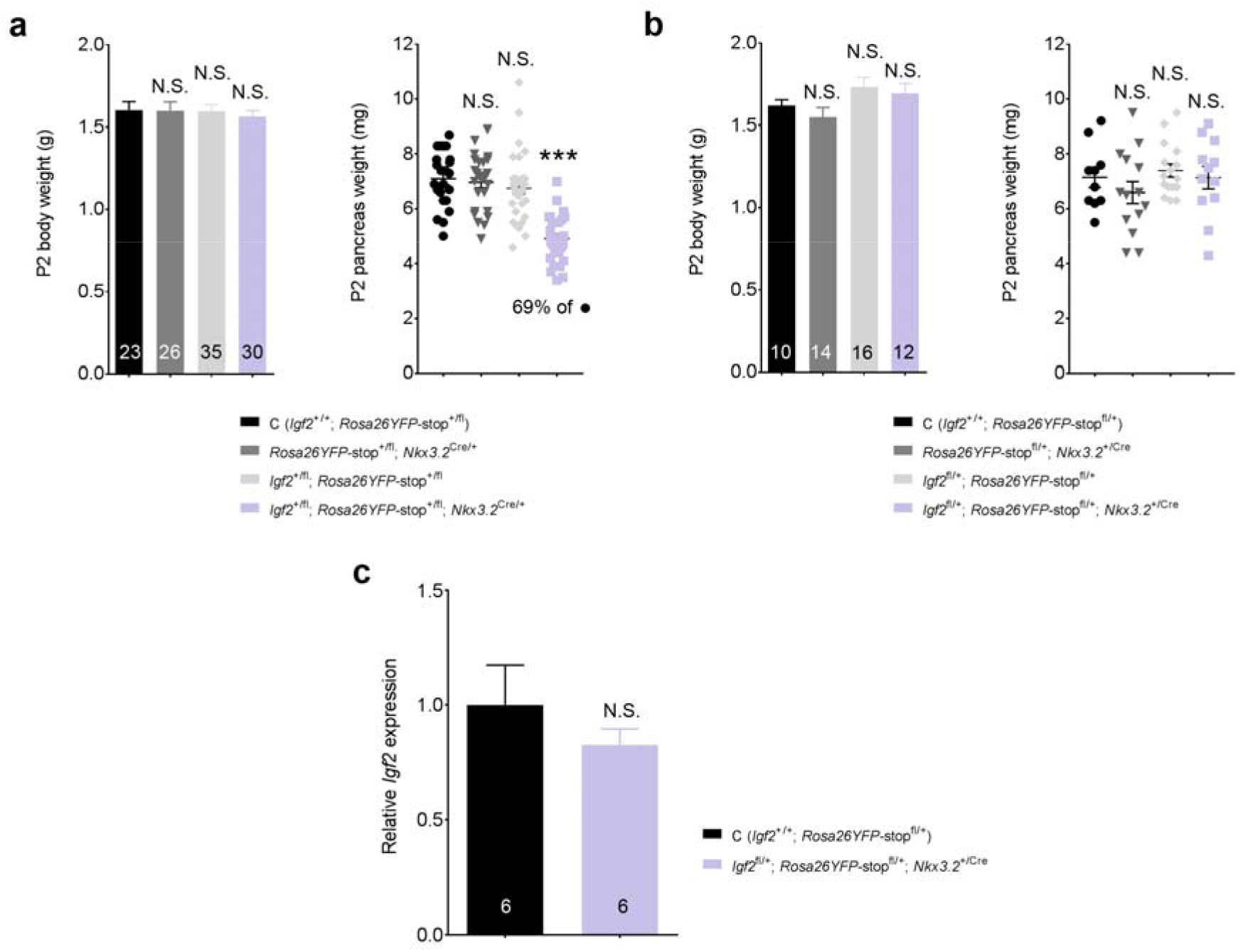
The presence of *Nkx3.2-Cre* or *Igf2* floxed alleles, or deletion of maternal *Igf2* alleles do not affect total body or pancreas weights at postnatal day 2 (P2). **(a)** body and pancreas weights in offspring obtained from a cross between heterozygous *Nkx3.2*-Cre females and heterozygous *Igf2* floxed males **(b)** body and pancreas weights in offspring obtained from a cross between heterozygous *Igf2* floxed females and heterozygous *Nkx3.2*-Cre males **(c)** *Igf2* mRNA levels measured by qRT-PCR in pancreases with a deletion of the maternal *Igf2* allele in the mesenchyme. Data is normalized to *Ppia* and shown relative to average *Igf2* levels in controls (C – *Igf2*^+/+^), set to 1. Data is shown as averages or individual values; error bars represent SEM. Numbers shown indicate numbers of animals for each genotype. Data was analysed using one-way ANOVA with Dunnett’s multiple comparison test against the control group for panels **(a)** and **(b)** and by unpaired Student’s t test in panel **(c)**; N.S. – non-significant; *** p<0.001.

**Supplementary Figure 6:**
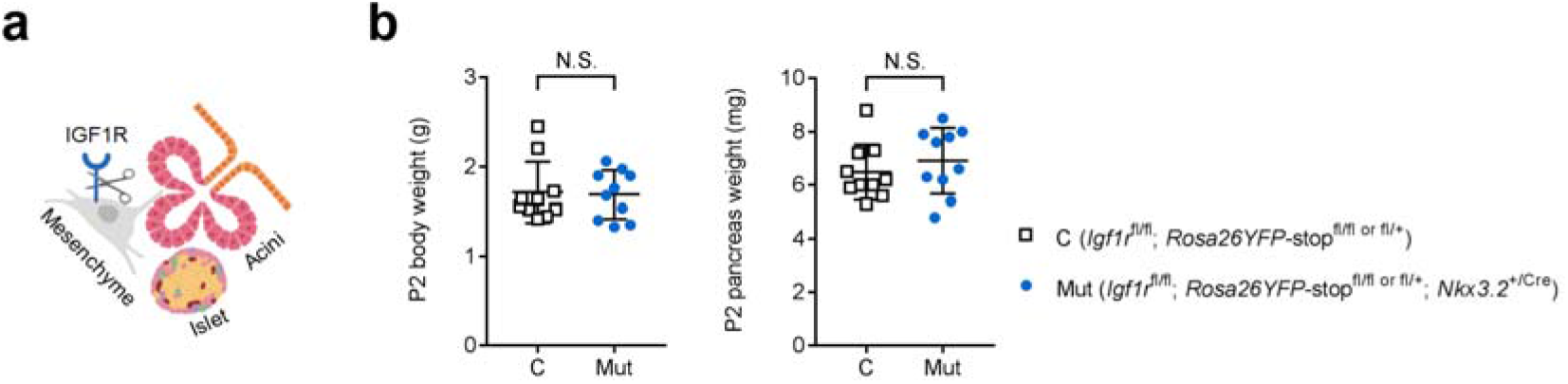
Normal pancreas growth upon mesenchyme-specific deletion of *Igf1r*. **(a)** Schematic representation of the conditional *Igf1r* deletion from the pancreatic mesenchyme. **(b)** Pancreas weights and total body weights are similar in mutants compared to littermate controls at postnatal day P2 (n=10 controls and n=10 mutants). Data is presented as individual values with averages ± SD; N.S. – non-significant by unpaired Student’s t tests.

**Supplementary Figure 7:**
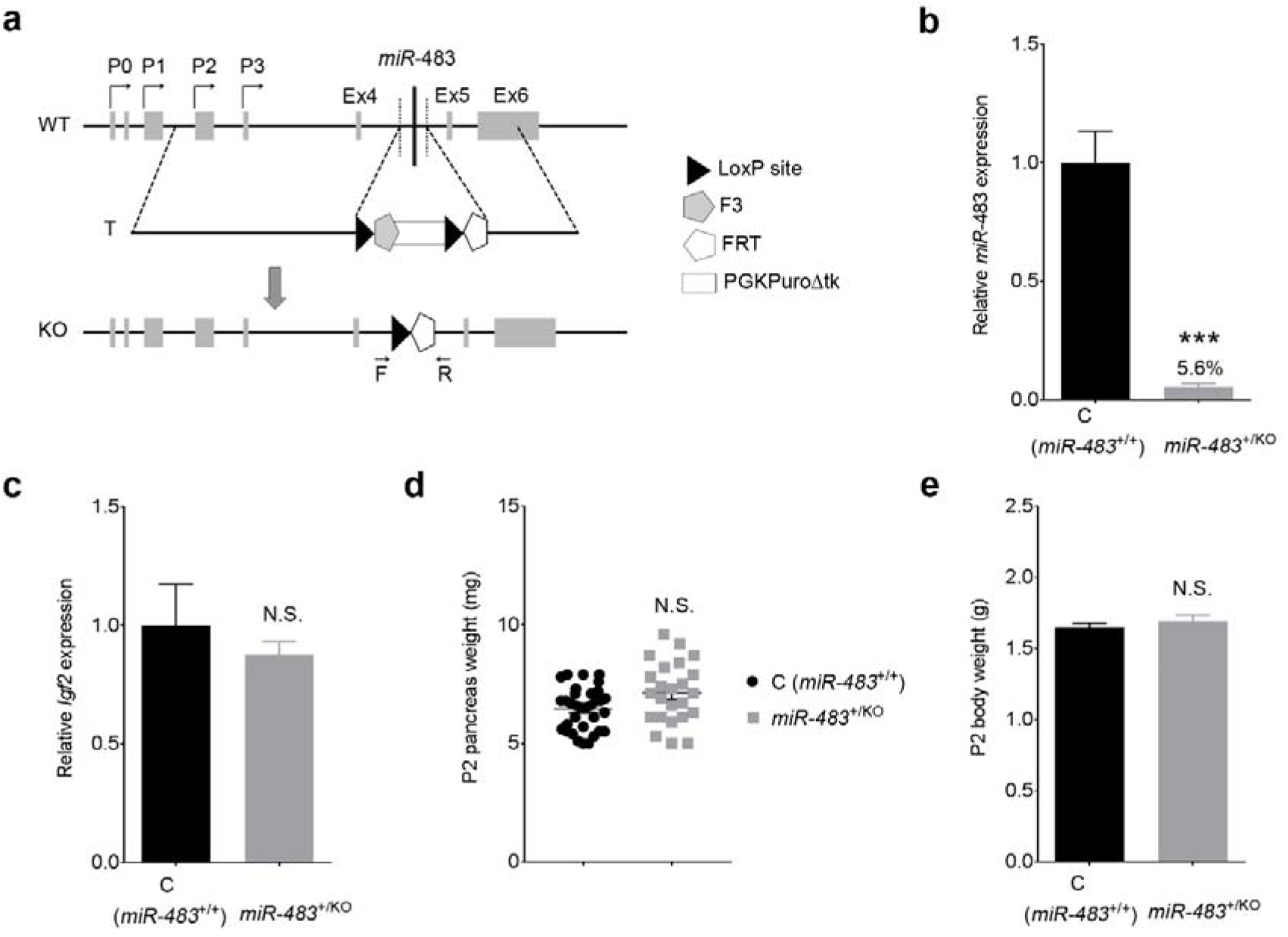
Paternally inherited *miR-483* deletion does not lead to reduction in pancreas size. **(a)** Gene targeting strategy to delete the intronic *Igf2 miR-483*. In brief, a targeting vector was used to replace the *miR-483* sequence in intron 4 of the *Igf2* gene with a PGKPuroΔtk selection cassette in ES cells. Removal of the selection marker was achieved by Cre-recombination between loxP sites. Germline transmitting *miR-483* KO chimeric mice were generated by ES cell injection into blastocysts (Sekita, Prosser, Zvetkova *et al.*, manuscript in preparation). (WT – wildtype allele, T – targeted allele, KO – knock-out allele, P0 – P3 are alternative *Igf2* promoters, Ex4 – Ex6 are the *Igf2* coding exons, F and R indicate the position of the forward and reverse primers used for PCR genotyping). **(b)** *miR-483* expression levels measured by qRT-PCR in postnatal day 2 (P2) whole pancreas from offspring of heterozygous *miR-483* KO males mated with wild-type females and shown relative (%) to littermate controls (C – *miR-483*^+/+^) set to 1. Expression data was normalized to *snoR-202* and *snoR-234* and is shown as average + SEM (n=10 C and n=10 KO). *** p<0.001 by unpaired Student’s *t* test. **(c)** *Igf2* mRNA expression is unaltered in *miR-483*^+/KO^. Data was normalized to *Ppia* and is shown relative to controls set to 1 (n=10 C and n=10 KO); error bars: SEM. N.S. – nonsignificant differences between genotypes by unpaired Student’s *t* test. Total pancreas weights **(d)** shown as individual values and population average ± SEM and corresponding body weights **(e)** at P2. Data is shown as averages (n=33 C and n=24 KO); error bars represent SEM; N.S. – non-significant differences by two-way ANOVA using genotype and litter as factors.

**Supplementary Figure 8:**
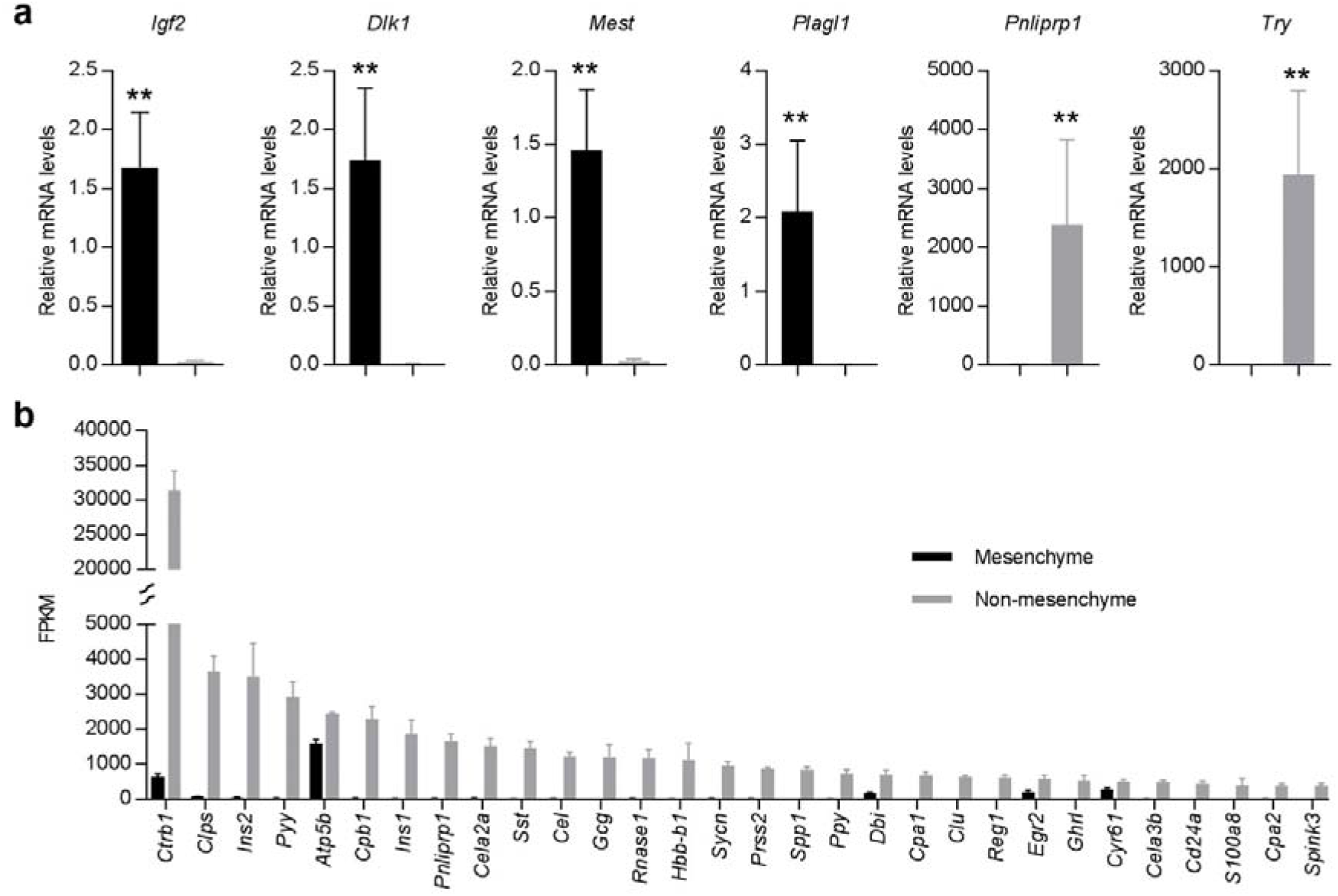
Gene expression analysis in pancreatic mesenchyme and non-mesenchyme cells at postnatal day 2 (P2). **(a)** Biological validation by qRT-PCR of differentially expressed genes between mesenchyme and non-mesenchyme, identified by RNA-seq (n=5-6 samples per group). Expression levels were normalized to *Ppia*. Data is shown are average values; error bars represent SEM; ** p<0.01 by Mann-Whitney tests. (b) Top 30 expressed genes with highest average FPKM values in pancreatic non-mesenchyme cells, and corresponding expression in mesenchyme (n=4; error bars represent SEM). Note that all 30 genes are significantly enriched in non-mesenchyme cells (>1.5 fold, FDR adjusted p value <0.05) compared to mesenchyme.

**Supplementary Figure 9:**
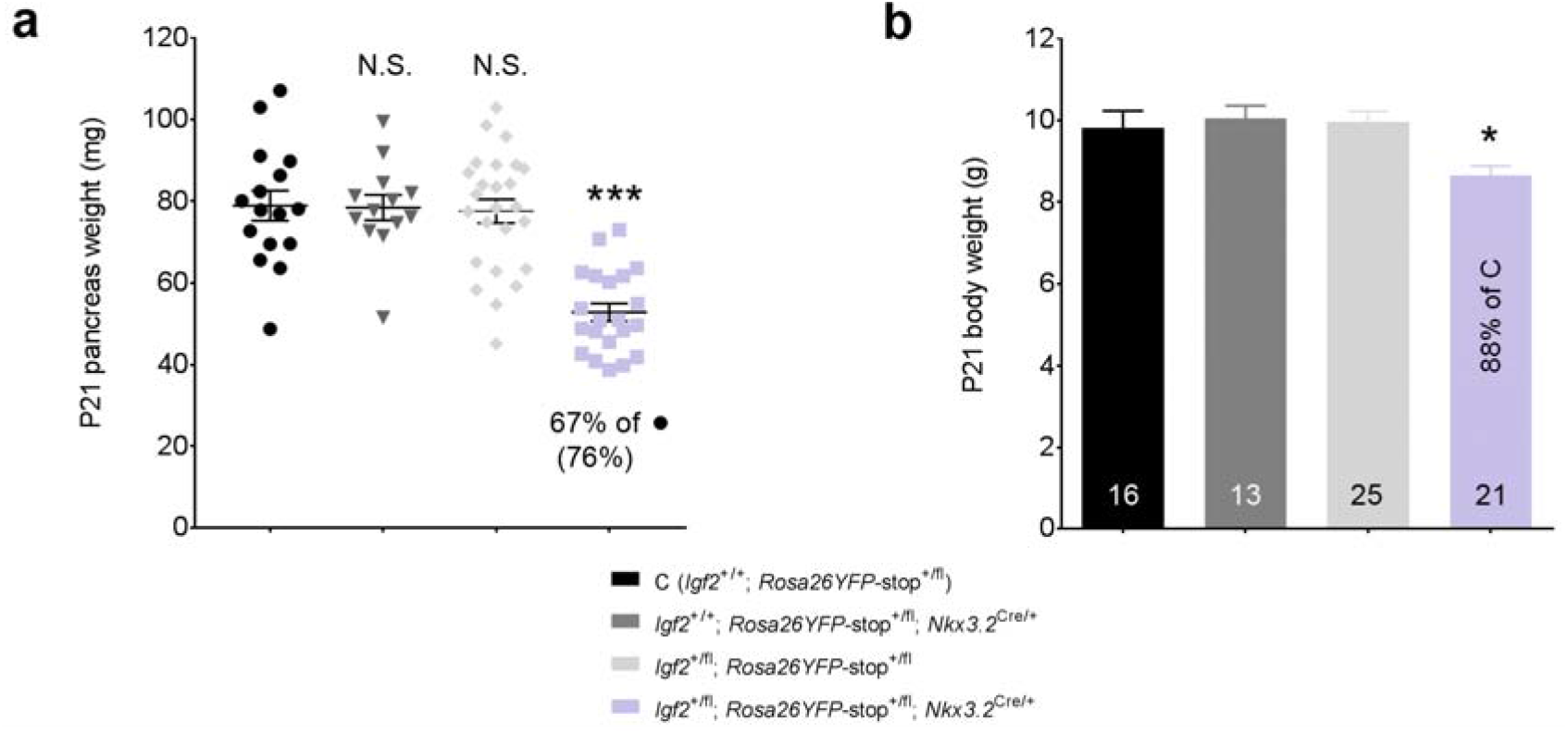
Deletion of paternal *Igf2* in pancreatic mesenchyme leads to reduced body and pancreas weights at weaning (P21). Total pancreas weight (a) and body weight (b) in offspring obtained from a cross between heterozygous *Nkx3.2*-Cre females and *Igf2* floxed males. The value within brackets in (a) shows % pancreas weight reduction after normalization to body weight. Data is shown as averages or individual values; error bars represent SEM. Numbers of mice for each genotype are shown. Data was analysed using one-way ANOVA with Dunnett’s multiple comparison tests against the control group (C – *Igf2*^+/+^); N.S. – non-significant; * p<0.05; *** p<0.001.

**Supplementary Figure 10:**
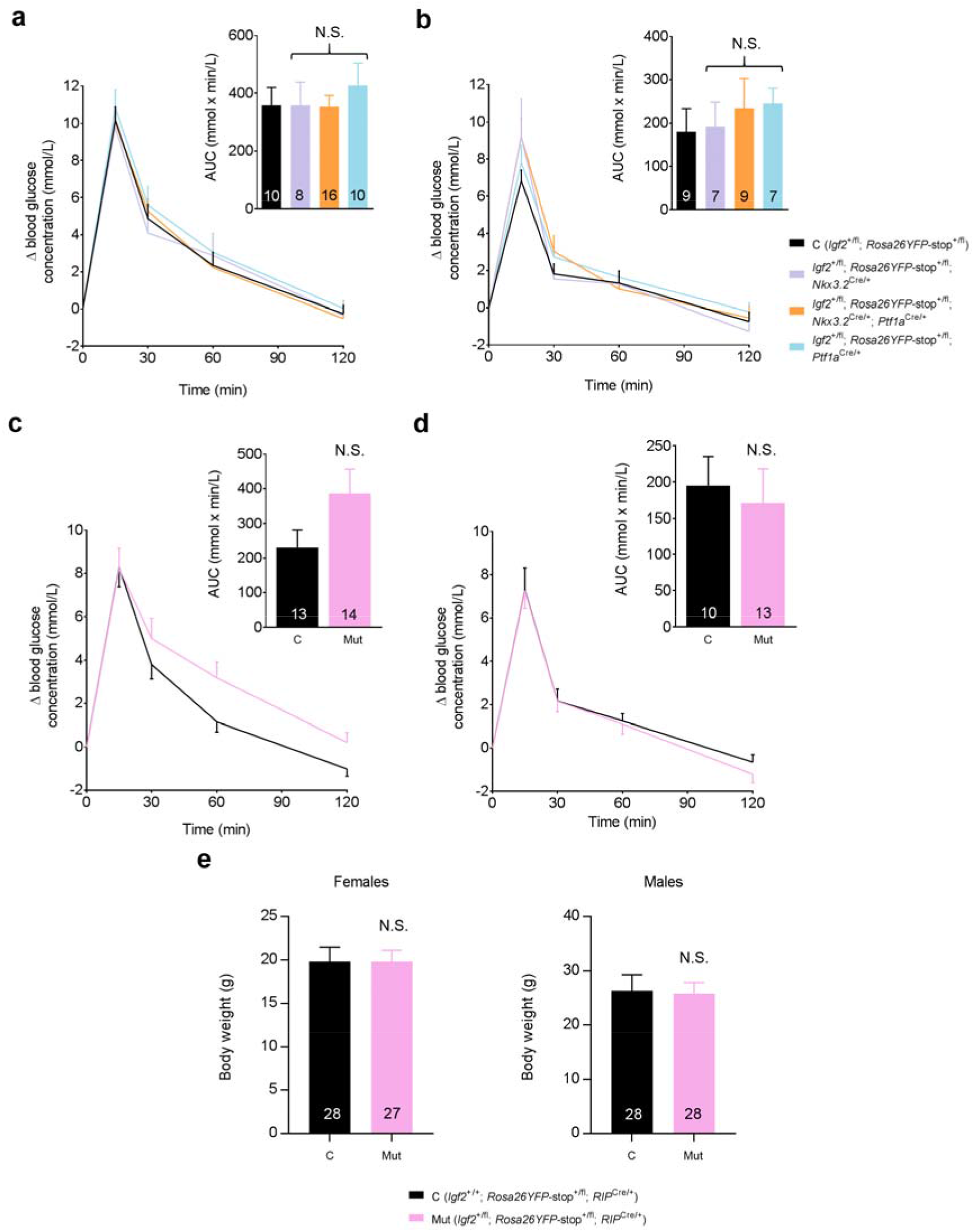
Glucose homeostasis and body weights in *Igf2* pancreas-cell type specific knockouts at 8 weeks of age. Oral glucose tolerance tests (OGTTs) in male mice (a) and (c), and female mice (b) and (d), with pancreas cell-type specific deletions of *Igf2* mediated by *Nkx3.2*-Cre: mesenchyme, *Ptf1a*-Cre: epithelium in (a) and (b) or *RIP*-Cre: beta-cells in (c) and (d). Panels (a) to (d) depict changes in blood glucose concentrations from basal pre-treatment values (Y axis) with time (X axis) after glucose administration. The insets indicate area under curve (AUC) calculated using the trapezoid rule. OGTT data was analysed statistically by repeated measures 1-way ANOVA with Dunnett’s multiple comparison tests against controls (C – *Igf2*^+/fl^) for panels (a) and (b) and by two-way ANOVA with Sidak’s multiple comparison tests for panels (c) and (d). AUC data was analysed statistically by 1-way ANOVA with Dunnett’s post-hoc tests for panels (a) and (b) and by unpaired Student’s t tests in panels (c) and (d). (e) Body weights in mice with beta-cell specific deletion of *Igf2*. Data was analysed using unpaired Student’s t tests. For all panels, data is shown as average values and the error bars represent SEM; N.S. – non-significant. Numbers of mice are shown within data columns (per genotype).

**Supplementary Table 1:**
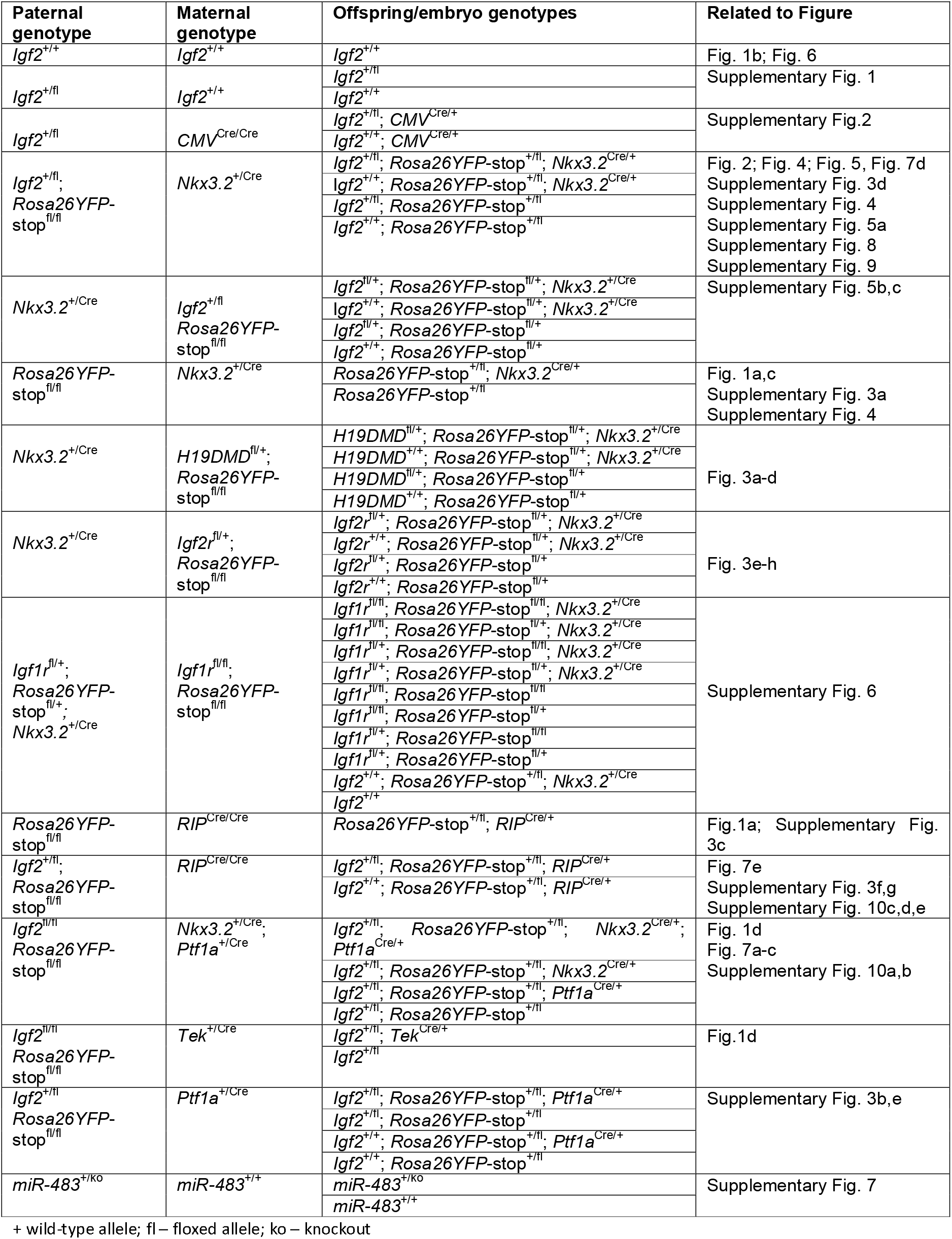
Mouse strains and crosses

**Supplementary Table 2:**
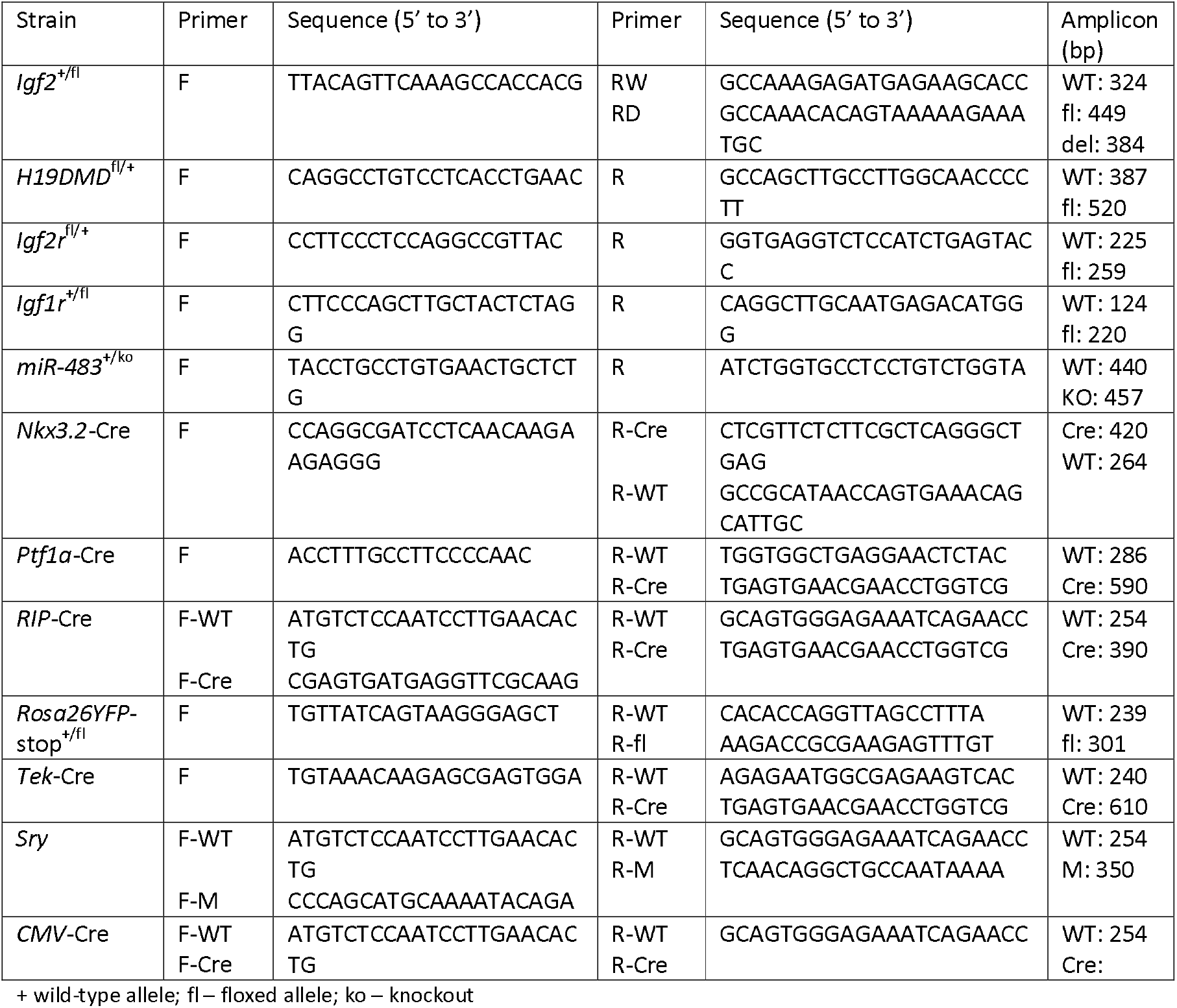
Primers used for genotyping by PCR

**Supplementary Table 3:** Conditions used for pancreas immunostaining

| Staining | Antigen retrieval | Blocking | Primary antibody | Secondary antibody |
| --- | --- | --- | --- | --- |
| Amylase | Autoclaving for 15 min at 121°C in citric acid buffer (10 mM citric acid, pH 6.0, 0.05% tween) | 5% Donkey serum (Sigma) | Rabbit polyclonal to amylase (1:100, Sigma A8273) overnight at 4°C | Donkey anti-rabbit (1:200, Jackson labs, Alexa Fluor 594) 1 hour at room temperature |
| YFP | Autoclaving for 15 min at 121°C in citric acid buffer (10 mM citric acid, pH 6.0, 0.05% tween) | 5% Donkey serum (Sigma) | Goat polyclonal to GFP (1:200, Abcam ab6673) overnight at 4°C | Donkey anti-goat (1:200, Jackson labs, Alexa Fluor 488) 1 hour at room temperature |
| IGF2 | Digestion with 1% pronase (Protease from Streptomyces griseus, Sigma Aldrich P6911) in 1xPBS for 10 min at 37°C | 15% Donkey serum (Sigma) | Goat anti-human IGF2 (1:50, R&D systems AF-292) overnight at 4°C | Donkey anti-goat (1:200, Jackson labs, Alexa Fluor 488) 1 hour at room temperature |
| Cytokeratin | Autoclaving for 15 min at 121°C in citric acid buffer (10 mM citric acid, pH 6.0, 0.05% tween) | 15% Donkey serum (Sigma) | Rabbit anti-cytokeratin (1:500, Dako) overnight at 4°C | Donkey anti-rabbit (1:200, Jackson labs, Alexa Fluor 594) 1 hour at room temperature |
| CD31 | Autoclaving for 15 min at 121°C in citric acid buffer (10 mM citric acid, pH 6.0, 0.05% tween) | 15% Donkey serum (Sigma) | Rabbit anti-CD31 (1:50, Abcam ab28364) overnight at 4°C | Donkey anti-rabbit (1:200, Jackson labs, Alexa Fluor 594) 1 hour at room temperature |
| Insulin | Not needed | 5% Rabbit serum (Dako) | Polyclonal guinea-pig anti-swine insulin antibody, (1:50, Dako A0564) 90 min at room temperature | Rabbit anti-guinea pig HRP coupled (1:100, Abcam ab6771) 1 hour at room temperature, then DAB for 2 min |

**Supplementary Table 4:**
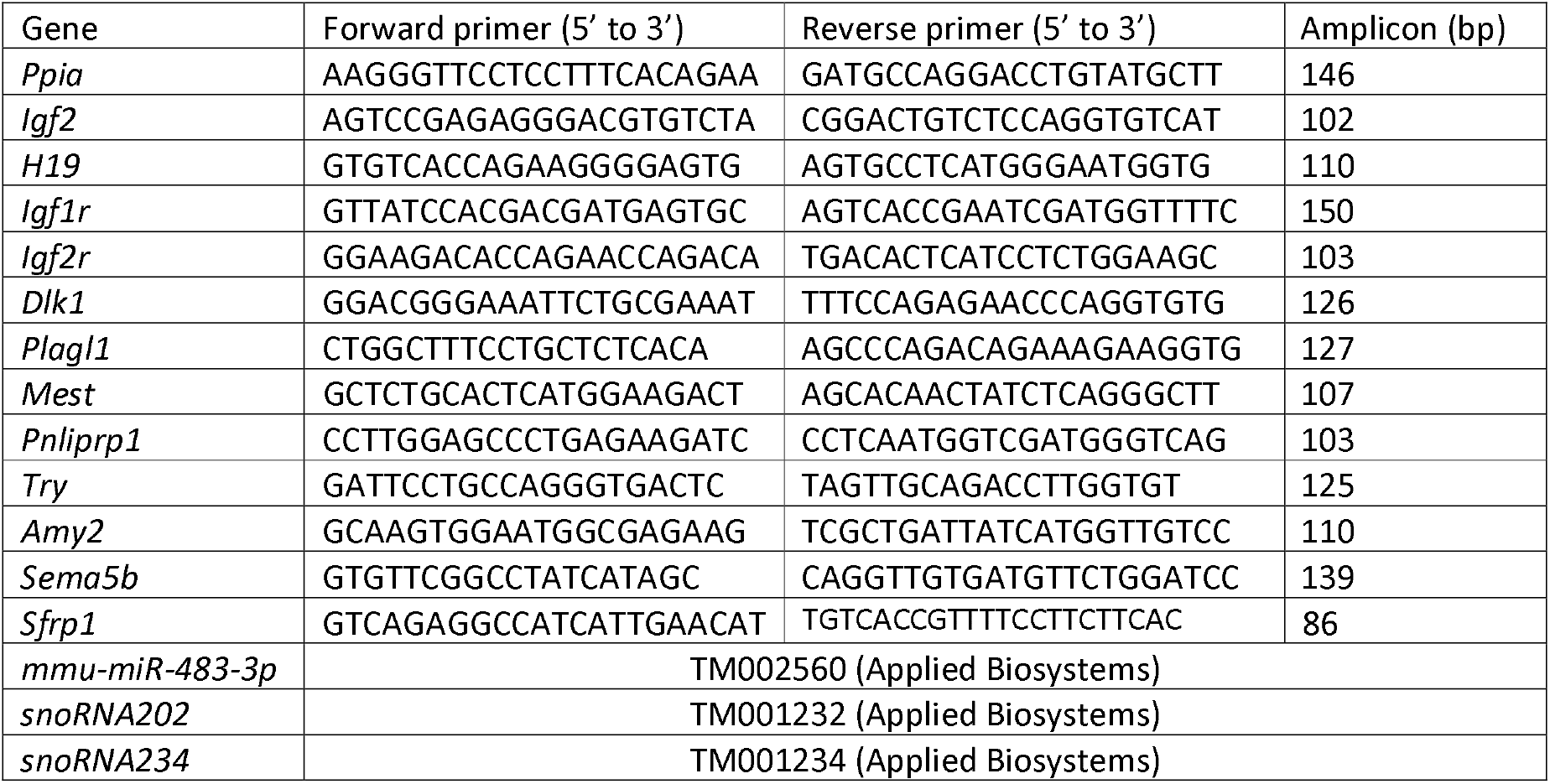
Primers/assays used for qRT-PCR

**Supplementary Data 1:** List of genes significantly enriched (fold change >1.5; FDR adjusted p value <0.05) in mesenchyme and non-mesenchyme pancreatic cells at P2 in wild-type mice (*Igf2*^+/+^; *Rosa26YFP*-stop^+/fl^; *Nkx3.2*^Cre/+^) and associated functional pathway analyses (DAVID).

**Supplementary Data 2:** List of differentially expressed genes (fold change >1.5; FDR adjusted p value <0.05) in non-mesenchymal cells between *Igf2*^+/fl^; *Rosa26YFP*-stop^+/fl^; *Nkx3.2*^Cre/+^ knockouts and *Igf2*^+/+^; *Rosa26YFP*-stop^+/fl^; *Nkx3.2*^Cre/+^ controls at P2, and associated functional pathway analyses (DAVID).

**Supplementary Data 3:** List of differentially expressed genes (fold change >1.5; FDR adjusted p value <0.05) in mesenchymal cells between *Igf2*^+/fl^; *Rosa26YFP*-stop^+/fl^; *Nkx3.2*^Cre/+^ knockouts and *Igf2*^+/+^; *Rosa26YFP*-stop^+/fl^; *Nkx3.2*^Cre/+^ controls at P2, and associated functional pathway analyses (DAVID).

